# Heat-shock pathway activation by TRC051384 protects spiral ganglion neurons from noise-induced hearing loss

**DOI:** 10.64898/2025.12.08.693048

**Authors:** Jintao Yu, Miguel A. Ramirez, Yi-Zhi Wang, Seby Edassery, Maxwell Shramuk, Mary Ann Cheatham, Mark A. Rutherford, Leah J Welty, Jeffrey N Savas

## Abstract

Noise-induced hearing loss (NIHL) is a major public health problem caused by damage to cochlear hair cells, synapses, and spiral ganglion neurons (SGNs). Since effective treatments are lacking, we investigated cellular stress responses induced by moderate and loud noise in a mouse model of cochlear synaptopathy. RNA sequencing and spatial transcriptomics revealed that noise exposure elicited a robust but transient upregulation of endoplasmic reticulum chaperones and proteasome subunits in SGNs and their supporting cells. To target this response, we administered TRC051384, a small-molecule activator of the heat shock transcription factor Hsf1, prior to noise exposure. TRC051384 crossed the blood–labyrinth barrier and reached the cochlea, induced heat shock protein gene expression, and restored ubiquitin–proteasome function in SGNs. Notably, TRC051384 treatment enhanced auditory brainstem response threshold recovery, preserved Wave I amplitudes, and maintained ribbon synapse density. These findings establish proteotoxic stress in SGNs as a key driver of NIHL and identify HSF1 activation as a promising therapeutic strategy.

## INTRODUCTION

Noise-induced hearing loss (NIHL) is a prevalent condition affecting greater than 5% of the global population, yet no effective therapies are currently available (1, 2). Exposure to harmful levels of noise is almost unavoidable in the industrialized world (3-5). NIHL is the result of a complex interplay of variables, including exposure duration, sound intensity and frequency, in addition to individual genetic and physiological differences (2). NIHL can result from exposure to an acute intense acoustic event, which causes immediate and irreversible damage to the inner ear. In contrast, chronic exposure to moderately loud sounds (typically above 85 dB SPL but below 100 dB SPL) often leads to subtle, progressive auditory deficits. Although these deficits may not be detected by standard audiometry, they are increasingly recognized as common and clinically significant and substantially increase the prevalence of NIHL (6, 7).

Loud noise adversely affects multiple cochlear structures, cell types, and cellular processes. For example, intense auditory stimulation imposes a high metabolic demand on the cochlea (8), leading to increased oxygen consumption, excessive free radical production, and cellular stress—all of which can contribute to cell death□(9-11). In addition, noise exposure has been shown to reduce both tectorial membrane calcium levels and cochlear blood flow□(10, 12), further exacerbating cochlear dysfunction. Hair cell (HC) death can occur through a variety of signaling pathways triggered by these stressors (13). Moreover, loud noise can lead to metabolic decompensation, characterized by nuclear and mitochondrial swelling, cytoplasmic vesiculation□(14), and ultimately apoptosis (15). Noise exposure can result in either permanent threshold shifts (PTS) due to irreversible damage, or temporary threshold shifts (TTS) that recover within a few weeks. The primary targets of noise-induced damage in the cochlea are supporting cells, cells of the lateral wall, the inner and outer hair cells (IHCs, OHCs) and their stereocilia bundles, and IHC ribbon synapses, onto auditory nerve fiber (ANF) terminals (16, 17). ANF terminals can physically detach from the presynaptic ribbons synapses of IHCs following noise exposure, but partial reattachment and functional recovery are sometimes observed (18-21). However, loud noise exposure can lead to the permanent loss of up to 50% of IHC ribbon synapses—even in cases where standard audiograms appear normal, a condition referred to as cochlear synaptopathy (22-25).

In our previous studies, we used quantitative, tandem mass spectrometry (MS)-based proteomic analysis to study cochlear extracts from mice immediately after they were exposed to loud noise. We discovered that noise levels sufficient to induce hearing loss caused a extensive protein accumulation in the cochlea (26). The level of the full proteasome complex, along with a broad spectrum of protein chaperones, was elevated. However, we mapped only a handful of elevated proteins to specific cochlear cell types. Furthermore, the interpretation of these results was complicated by the fact that the sound pressure level (SPL) of the noise exposures caused compound threshold shifts, including both a temporary and a permanent component. Making it difficult to distinguish adaptive cellular responses from ineffective compensation for structural damage. In additional MS-based proteomic analyses, we found that the translational machinery (i.e., ribosomal protein sub-units) was selectively elevated during at two weeks post-noise exposure, thereby suggesting that loud noise (94 dB SPL, 0.5 hour, 8-16 kHz) triggered a burst of protein turnover that may play a key role in restoring hearing sensitivity (27).

Building on these findings, the present study primarily investigated a mouse model of cochlear synaptopathy. After validating the noise exposure paradigm, we performed discovery-based bulk RNA sequencing to characterize the timeline of gene expression induced by noise exposure causing cochlear synaptopathy. Notably, a coordinated peak in gene expression was observed four hours after the two-hour exposure to 100 dB SPL. Pathway enrichment analysis of the significantly elevated genes revealed a striking enrichment of the endoplasmic reticulum (ER) chaperone complex, emerging as the most significantly overrepresented term by a wide margin. This was driven by the increased gene expression of several protein chaperones, including *Hsp90b1* and *Hspa5* encoding Endoplasmin and BiP respectively. Spatial transcriptomics revealed that both spiral ganglion neurons (SGNs) and their associated glia cells exhibited elevated expression of genes encoding proteasome subunits. Next, we examined whether pre-activating the heat shock response, a pathway activated following stressors that cause protein misfolding, could protect against noise-induced damage. To do so, we pharmacologically activated the transcription factor Hsf1 prior to noise exposure, using the previously identified small molecule TRC051384 (28). Using a ubiquitin-proteasome system (UPS) activity reporter mouse line, we observed that cellular stress in response to synaptopathic noise exposure was primarily localized to the spiral ganglion. Notably, pre-treatment with TRC051384 mitigated the noise-induiced accumulation of a GFP-based reporter, indicating enhanced proteostasis and improved protein degradation capacity following noise-induced stress with activation of the heat shock response. Finally, we found that the prophylactic administration of TRC051384 not only improved the recovery of auditory brainstem responses (ABR) Wave I amplitudes but also conserved synaptic density.

## RESULTS

### Moderate noise exposure causes cochlear synaptopathy in 16-week-old CBA/CaJ mice

ABRs are sound-evoked potentials that can be used to determine abnormalities or deviations in hearing sensitivity. The first wave in the ABR represents the synchronous auditory nerve discharge at the onset of sound, proportionate to activity of the cochlear ribbon synapses between IHCs and SGNs. Initial experiments aimed to replicate the previously reported “hidden hearing loss” phenotype in CBA/CaJ mice (22). Thus, we exposed 8- and 16-week-old CBA/CaJ mice to 8-16 kHz octave-band noise at 100 dB SPL for 2 hours. ABRs and distortion product otoacoustic emissions (DPOAEs) were measured at baseline and at 1-, 3-, and 14-days after noise exposure. Both the 8- and 16-week-old mice exhibited significantly elevated ABR thresholds (i.e., the lowest tone level necessary to generate the ABR waveform) post noise The largest threshold shift, 40 dB, was observed for tones of 32 kHz. Elevated thresholds persisted at frequencies above 20 kHz for at least 3 days after noise (DAN). As expected, ABR thresholds recovered to near baseline levels by 14 DAN in 16-week-old but not 8-week-old CBA/CaJ mice **(Figure 1A and S1A)**. In 16-week-old-mice, Wave I amplitudes in response to 32 kHz tones remained significantly reduced (∼50%) at 14 DAN **(Figure 1B, C)**. This lasting reduction in ABR Wave I amplitude following a temporary threshold shift (TTS) is consistent with permanent cochlear synaptopathy. In 8-week-old mice, Wave I amplitudes failed to recover to pre-noise levels at frequencies greater than 16 kHz **(Figure S1B-D)**.

**Figure 1.**
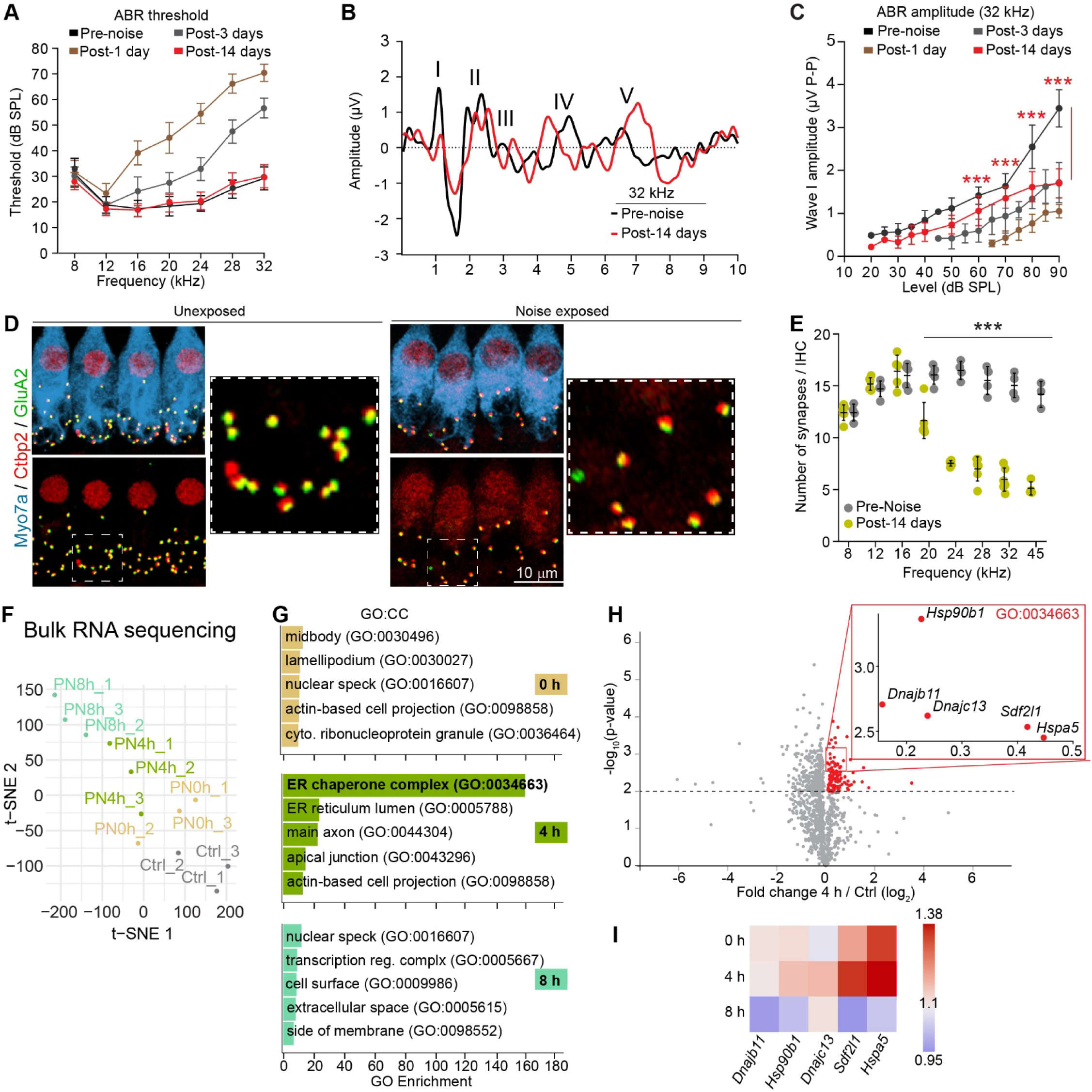
Exposure to 100 dB SPL noise for two hours reproduces the hidden hearing loss phenotype in male mice and induces a robust wave of ER chaperone gene expression in the 16-week-old CBA/CaJ mouse cochlea. **(A)** ABR thresholds in 16-week-old male CBA/CaJ mice show no significant difference 14 days after noise (DAN) compared to before noise (BN). **(B)** Representative ABR traces at 90 dB SPL BN and 14 DAN at 32 kHz. **(C)** Wave I amplitudes at 32 kHz with increasing intensities. Wave I amplitudes failed to recover to baseline levels by 14 DAN across the 60-90 dB SPL stimulus range compared to BN (60 dB SPL estimated difference = -0.748 adjusted 95% CI [-1.21, -0.29], 70 dB SPL estimated difference = -0.953 adjusted 95% CI [-1.41, -0.496], 80 dB SPL estimated difference = -0.724 adjusted 95% CI [-1.18, -0.267], 90 dB SPL estimated difference = -1.65 adjusted 95% CI [-2.11, -1.2]. **(D)** Representative images from 32 kHz regions of the cochlea immunostained with antibodies for CtBP2 (red) and GluA2 (green) in an unexposed control cochlea and a noise-exposed cochlea 14 DAN. Scale bar = 10 µm. **(E)** Quantification of overlapping CtBP2 and GluA2 puncta from (D) revealed a dramatic loss of synapses above 20 kHz 14 DAN (average synapses per IHC from 20-45 kHz = 7.45 ± 2.53) compared to a control unexposed cochlea (average synapses per IHC from 20-45 kHz = 15.46 ± 0.89). **(F)** □t-SNE plot of bulk-cochlear RNA-sequencing profiles from individual mice collected immediately after the 100□dB□SPL noise exposure, 4□h post noise (PN)-exposure, 8□h PN-exposure, and from unexposed controls, demonstrating that biological replicates cluster according to exposure status and recovery time. **(G)** Gene Ontology (GO) cellular component (CC) enrichment analysis of the genes with significantly elevated expression 4 hours, or 8 hours post exposure relative to unexposed controls. The most significantly enriched term was “endoplasmic reticulum chaperone complex (GO:0034663)” in the 4-hour post exposure group. **(H)** Volcano plot depicting noise-induced changes in cochlear gene expression in the 4-hour post exposure group relative to unexposed controls. Genes with significantly elevated fold change (Student’s t-test p value < .01) are red. Insert represents the 5 genes associated with the GO:CC term endoplasmic reticulum chaperone complex (GO:0034663). **(I)** Heat map depicting the gene expression of *Hsp90b1*, *Dnajb11*, *Dnajc13*, *Sdf2l1*, and *Hspa5* revealing elevated expression immediately following noise exposure and peaking at 4 hours post-exposure relative to unexposed control mice. (A) Data are represented as mean ± SD, *** = p value < .001 by one-way ANOVA determined significance with Tukey’s post hoc correction. (C) Differences were estimated by linear mixed modeling. (E) Data are represented as mean ± SD. *** = p value < .001 by one-way ANOVA with Tukey post hoc correction. (A-C) N = 6 male mice, (D-E) N = 4 unexposed, 6 exposed male mice (1 cochlea per mouse). (F-I) N = 4 male mice per group (2 cochleae per mouse).

DPOAEs are acoustic responses generated by vibrations along the basilar membrane when two pure tones stimulate the cochlea. These emissions reflect activity of the cochlear OHCs. Elevation of DPOAE thresholds indicates dysfunction of the OHCs, upstream of auditory nerve excitation by ribbon synapses. DPOAE thresholds were elevated at 1 DAN in both ages of mice **(Figure S1E-K)**. In contrast to ABRs, DPOAE thresholds and amplitudes fully recovered at all tested frequencies by 14 DANs, indicating that OHCs were not permanently damaged by these noise exposures in either group of mice.

In the absence of threshold shifts, a reduction of ABR wave I amplitude indicates damage or loss of cochlear ribbon synapses. To verify this finding, we examined immunostained whole mounts of the organ of Corti with antibodies against CtBP2 and GluA2, which stain pre-synaptic ribbons and post-synaptic receptors, respectively **(Figure 1D)**. Juxtaposition of these two markers indicates a synapse. Consistent with our ABR data at 14 DAN, we found significant reductions in cochlear ribbon synapse density at all frequencies above 20 kHz **(Figure 1E)**. In the 32 kHz region, the average number of ribbon synapses per IHC was 15.0 ± 1.1 in cochlea from age matched unexposed CBA/CaJ mice and 6.0 ± 1.1 at 14 DAN. Similar results were obtained when separately counting either pre-synaptic ribbons or post-synaptic puncta **(Figure S1L,M)**. Together, our results confirm that this level of noise exposure produced a TTS while inducing cochlear synaptopathy that is thought to be permanent in CBA/CaJ mice.

### A wave of protein chaperone gene expression peaking four hours after synaptopathic noise

We next sought to identify cellular pathways engaged acutely after synaptopathic noise exposure. We repeated our noise exposure paradigm in a new cohort of 16-week-old mice and harvested total cochlear RNA at 0-, 4-, or 8-hrs after noise. Bulk cochlear gene expression was measured using paired-end RNA sequencing (RNA-Seq) analysis (29). We obtained greater than 40 million mapped reads at a total mapping rate of > 96.91% from each of the three biological replicates per group. To investigate the reproducibility of our transcriptomic analysis, we assessed the correlation of gene expression patterns between the individual biological datasets using t-distributed stochastic neighbor embedding (t-SNE). Notably, the biological replicates clustered according to the duration after exposure **(Figure 1F)**.

We assessed changes in gene expression at each time point relative to unexposed controls, identifying 158, 118, and 74 genes with significantly elevated levels at 0-, 4-, and 8-hours after noise, respectively **(Table S1)**. To identify the pathways engaged in response to this noise exposure paradigm, we performed Gene Ontology (GO) cellular component (CC) enrichment analysis using DAVID (30). Among all three replicates, the most significantly enriched term was the GO:CC category “endoplasmic reticulum chaperone complex (GO:0034663),” which was detected only at 4 hr post-noise **(Figure 1G and Table S2)**. This 159.7-fold enrichment of the endoplasmic reticulum chaperone complex was 7-fold greater than the second most enriched GO:CC term. To visualize noise-induced changes to cochlear gene expression, we graphed the fold changes observed 4 hours after noise exposure relative to unexposed controls on a volcano plot **(Figure 1H)**. The five genes in our dataset associated with GO:0034663 were *Hsp90b1*, *Hspa5, Dnajb11*, *Dnajc13*, and *Sdf2l1*. Close inspection of gene expression time-course revealed a slight elevation immediately after noise, peaking 4 hours after noise, and resolving to near baseline levels by 8 hours **(Figure 1I)**. The protein products of these genes play well known and important roles in regulating protein quality control. *Hsp90b1* (i.e., endoplasmin) is an ER-resident chaperone, functioning in protein folding and quality control. *Hspa5* (i.e., BiP) is the master chaperone in the ER and is central to the unfolded protein response (UPR). *Dnajb11* (i.e., ERdj3) is an ER co-chaperone that works with BiP to assist in protein folding. *Dnajc13* (i.e., RME-8) a co-chaperone involved in endosomal trafficking and implicated in membrane dynamics. *Sdf2l1* encodes an ER stress-induced protein, also participating in protein folding.

To investigate whether expression of these genes is upregulated in all cells or in a select group of cells following exposure, we utilized RNAScope based *in situ* hybridization to visualize patterns of *Hsp90b1*, *Hspa5* and *Sdf2l1* gene expression post noise exposure. Expression of all three genes was markedly elevated in the spiral ganglion, IHCs, and OHCs at X hrs post noise **(Figure S2)**. Interestingly *Hsp90b1* expression also appeared to increase around the spiral limbus. Thus, RNAScope in situ hybridization revealed that the primary sensorineural cells contributed to the elevated expression of genes involved in ER protein quality control in the cochlea.

### GeoMx spatial transcriptomics reveals elevated proteasome subunit gene expression in SGNs and support cells following noise exposure

To further investigate the transcriptional response of cochlear cells to noise exposure, we performed GeoMx spatial transcriptomic profiling on cochlear sections from mice exposed to 100 dB SPL for 2 hours, the cochleae were harvested 4 hours post-exposure, alongside mock-exposed controls. Immunofluorescence imaging identified NF200-positive SGNs, which were segmented into regions of interest (ROIs) for GeoMx spatial transcriptomic analysis. Each ROI was further subdivided into a neuronal mask (i.e., NF200 positive) and a supporting cell mask (i.e., NF200 negative) to distinguish expression profiles between SGNs and ganglionic supporting cells **(Figure 2A-F and Table S3)**. Dimensionality reduction using t-SNE revealed that RNA expression profiles clustered primarily by cell type and, to a lesser extent, by noise exposure condition **(Figure 2G)**. In SGNs, noise exposure induced substantial upregulation of genes encoding the proteostasis network, particularly multiple subunits of the proteasome **(Figure 2H)**. Consistently, GO enrichment analysis of the upregulated genes identified the proteasome complex as the only cellular component category significantly enriched **(Figure 2I and Table S4)**. Similarly, support cells also exhibited noise-induced upregulation of mRNAs encoding proteostasis factors, with GO enrichment analysis identifying significant enrichment in the proteasome complex and nucleus **(Figure 2J-K and Table S4)**. Taken together, these data suggest that both SGNs and their supporting cells activate proteasome-associated transcriptional programs in response to synaptopathic noise.

**Figure 2.**
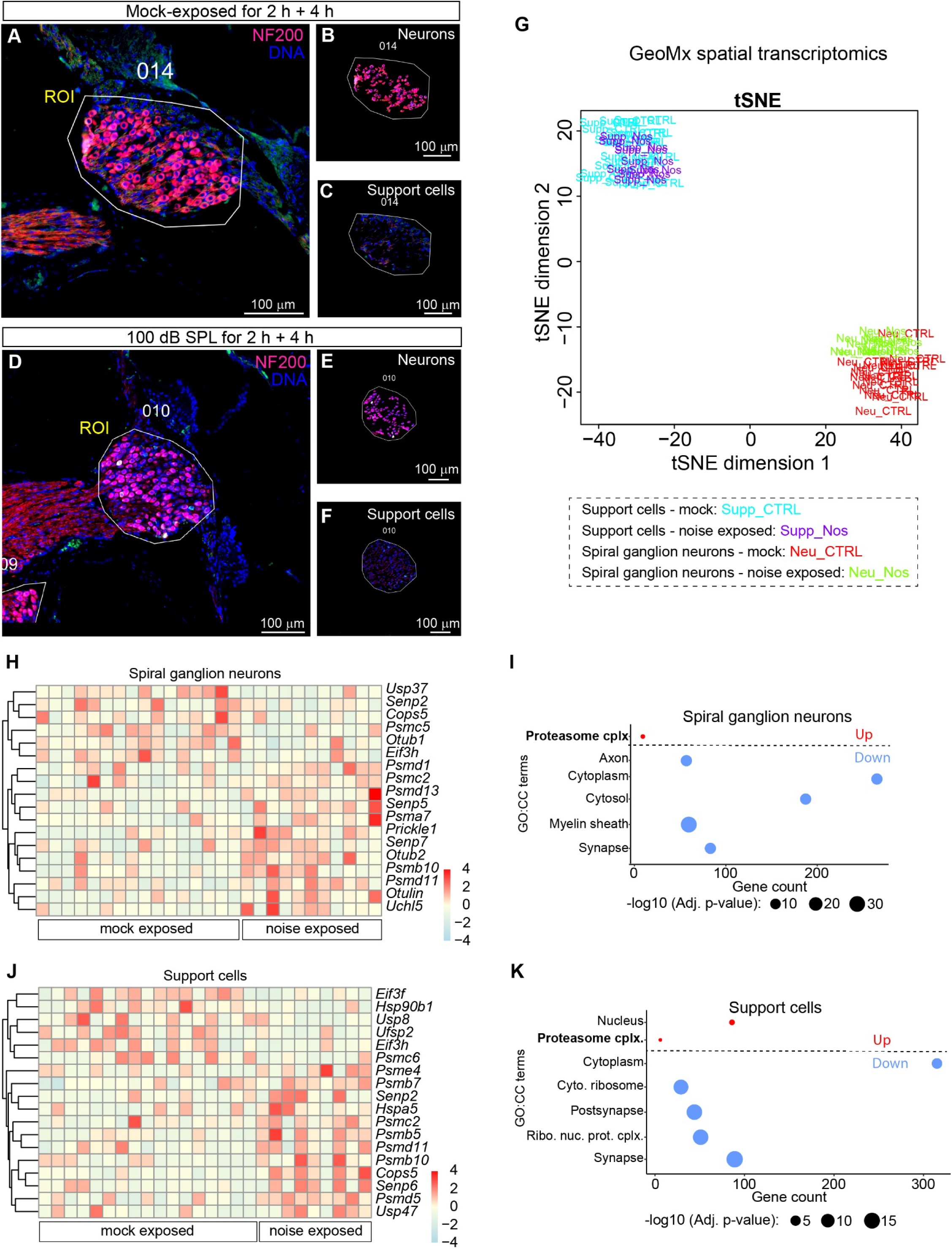
GeoMx spatial transcriptomics reveal that noise exposure selectively upregulates genes encoding multiple proteasome subunits in spiral ganglion neurons and associated support cells. **(A–F)** Representative GeoMx immunofluorescent cochlear images from a mock control mouse and an experimental mouse exposed to 100 dB SPL for 2 hours and harvested 4□h later. NF200-positive neuronal somata are in magenta; nuclei (Hoechst) are blue. Regions of interest (ROI; white outline) encompassing the spiral ganglion were segmented into a neuronal mask (B, NF200 +ive) and a support-cell mask (C, NF200 +ive) for probe collection. **(G)** t-SNE plot of RNA-sequencing profiles from individual spiral ganglion neuron ROIs, showing that biological replicates cluster primarily by cell type and, to a lesser extent, by noise-exposure condition. Supp_CTRL (blue) = supporting cells from mice not exposed to noise, Supp_Nos (purple) = supporting cells from mice exposed to noise, Neu_CTRL (red) = SGNs from mice not exposed to noise, Neu_Nos (green) = SGNs from mice exposed to noise, **(H)** Heat map showing genes encoding proteostasis-related proteins with significantly altered mRNA levels in spiral ganglion neurons (SGNs). Each row represents a gene, and each column represents a region of interest (ROI). Z-scored expression values are shown, with red indicating upregulation and blue indicating downregulation in response to noise exposure. **(I)** Database for Annotation, Visualization, and Integrated Discovery (DAVID) enrichment analysis of the upl11regulated genes in SGNs shows only one term that surpasses the significance threshold: the proteasome complex (KW-0647∼Proteasome). **(J)** Heat map showing genes encoding proteostasis-related proteins with significantly altered mRNA levels in support cells. Each row represents a gene, and each column represents a region of interest (ROI). Z-scored expression values are shown, with red indicating upregulation and blue indicating downregulation in response to noise exposure. **(K)** DAVID enrichment analysis of the upl11regulated genes shows only two terms surpassing the significance threshold: the proteasome complex (KW-0647∼Proteasome) and nucleus (KW-0539∼Nucleus). N = 11 (exposed) and 16 (control) ROIs for SGNs, N = 9 (exposed) and 17 (control) ROIs for support cells from N = 5 P60-65 mice of mixed sex per condition.

### TRC051384 induces the heat shock response

TRC051384 is a small-molecule activator that robustly induces the cellular heat shock response. It belongs to the 2-propen-1-one chemical class and functions as a potent inducer of heat shock protein (Hsp) expression via activation of heat shock factor 1 (Hsf1)(28). To verify that TRC051384 effectively engages canonical Hsf1 signaling, we examined the transcriptional activation of Hspa1a, a well-characterized Hsf1 target gene. HEK293T cells were treated with either vehicle control or TRC051384, and transcript levels were quantified by qPCR. TRC051384 treatment led to a dramatic (>20-fold) increase in Hsp1a expression at the 4-hour timepoint compared to vehicle-treated cells, while *Hsf1* mRNA levels remained unchanged, consistent with the mechanism of post-translational activation of Hsf1 **(Figure 3A)**. To determine whether this transcriptional response could be reproduced in inner ear cells, we performed qPCR on cochlear organotypic cultures derived from postnatal day 4 (P4) rat pups. Remarkably, a similar expression pattern was observed, with *Hspa1a* showing its strongest induction at the 4-hour mark **(Figure 3B)**, albeit with a smaller fold change than in HEK293T cells. This attenuated response in cochlear tissue may reflect cell type–specific differences in basal Hsf1 activity, accessibility of the *Hspa1a* promoter, or pharmacokinetic factors influencing compound uptake. Together, these findings confirm that TRC051384 is a potent pharmacological activator of the heat shock response in both immortalized human cells and *ex vivo* mammalian cochlear tissue, reinforcing its utility as a molecular tool for inducing Hsf1-mediated stress protection pathways.

**Figure 3.**
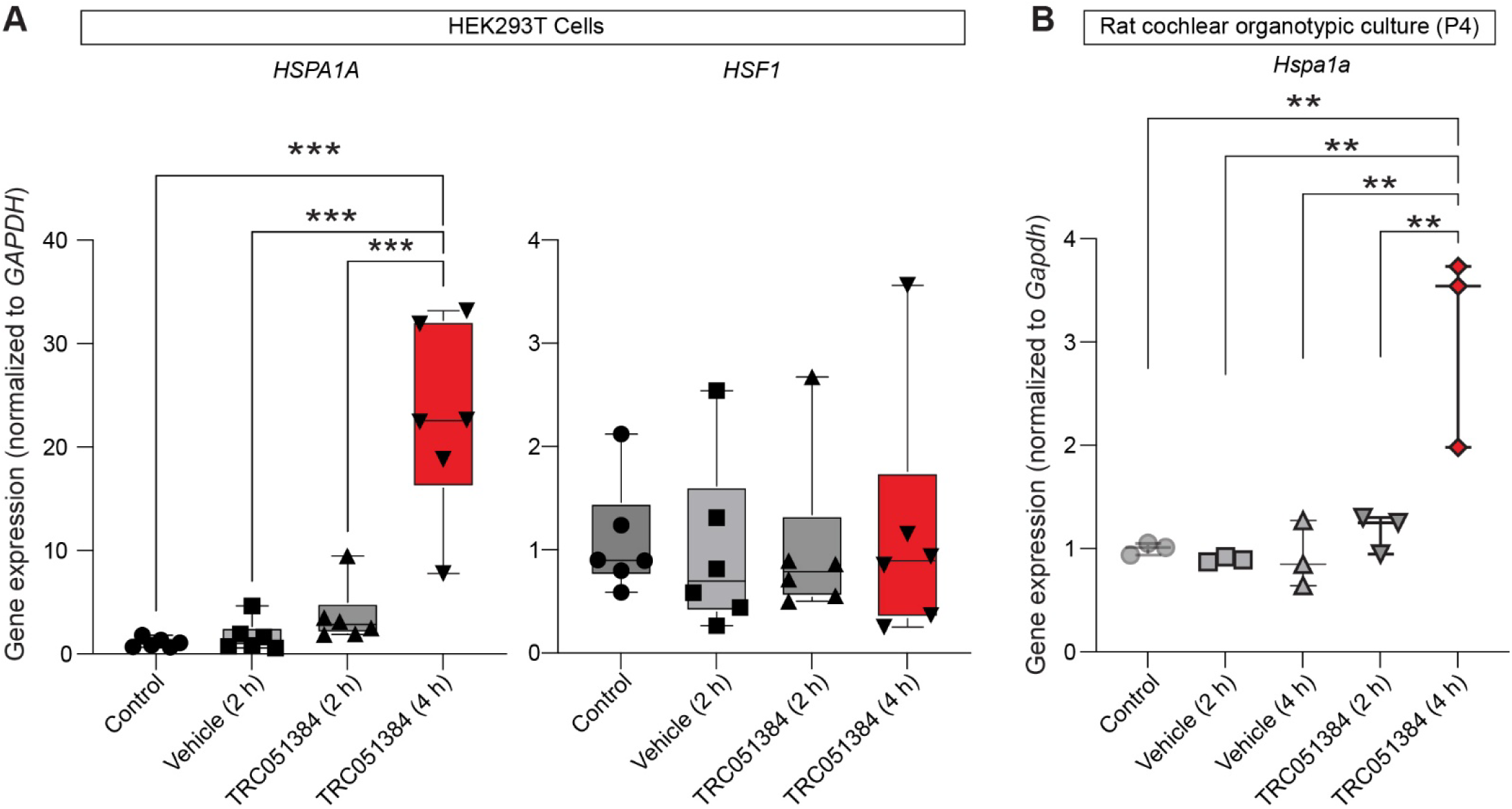
TRC051384 selectively enhances *HSPALA* gene expression without altering *HSF1* expression in HEK293T cells and cochlear organotypic cultures. **(A)** QPCR analysis of RNA from HEK293T cells treated with TRC051384 or with a DMSO vehicle, relative to untreated controls. Cells were untreated (Control) or incubated with DMSO for 2 hours or TRC051384 for 2 or 4 hours prior to RNA collection. *HSPA1A* (Heat Shock Protein Family A (Hsp70) Member 1A) expression was significantly increased by TRC051384 treatment, with levels rising further at 4 hours relative to DMSO and untreated controls (Control: 1.07 ± 0.45; DMSO: 1.73 ± 1.52; 2 h TRC051384: 3.75 ± 2.88; 4 h TRC051384: 20.10 ± 11.5). *HSF1* mRNA levels remained unchanged across all conditions over the 4-hour period (Control: 1.09 ± 0.55; DMSO: 0.99 ± 0.84; 2 h TRC051384: 1.06 ± 0.91; 4 h TRC051384: 1.18 ± 1.20). N = 6 independent biological replicates per condition. **(B)** QPCR analysis of RNA collected from P4 rat cochlear organotypic cultures at DIV3. Cultures were untreated (Control) or incubated with DMSO for 2 hours or TRC051384 for 2 or 4 hours prior to RNA collection. *Hspa1a* (i.e., Heat Shock Protein Family A (Hsp70) Member 1A) gene expression significantly increased by 4 hours relative to DMSO and untreated controls (Control: 1.00 ± 0.06, DMSO 2 hour: 0.89 ± 0.02, DMSO 4 hour: 0.92 ± .32, 2Hour TRC051384: 1.17 ± 0.19, 4Hour TRC051384: 3.08 ± 0.96). N = 3 independent cultures per condition. For (A + B) Data are represented as mean ± SD, ** = p value < .01 and ***p < .001 by one-way ANOVA determined significance with Tukey’s post hoc correction.

To evaluate the pharmacokinetic accessibility of TRC051384 to the inner ear, we examined its ability to penetrate the cochlea after systemic administration. Mice received intraperitoneal injections of TRC051384 (60 mg/kg). Then, the cochleae were dissected and processed to solubilize small molecules. We analyzed TRC051384 alone and in the resulting cochlear extracts using MS1 targeted selected ion monitoring (tSIM) (31), to sensitively and selectively quantify TRC051384 based on its molecular mass (465.23 Da; +2 charge state = m/z 233.6261). Evidence of drug penetration was clearly observed: extracts from the cochleae of TRC051384-treated mice displayed a strong peak at m/z 233.62, which eluted at 30–31 minutes. This signal was entirely absent from the extracts of vehicle-treated controls (**Figure S3A–B**). To further confirm the identity of the compound within the complex biochemical environment of the cochlea, we performed a fragment ion search by coupling tSIM with data-dependent MS2 acquisition. This analysis revealed numerous diagnostic TRC051384 fragment ions, unambiguously verifying its presence in cochlear tissue (**Figures S3C–D**). These findings demonstrate that TRC051384 readily crosses the blood-labyrinth barrier and accumulates in the cochlea at quantifiable levels, establishing its pharmacological accessibility to inner ear tissues. This property provides a strong foundation for exploring TRC051384’s ability to target heat shock pathways as a potential therapeutic strategy for auditory protection and repair.

### TRC051384 alleviates noise-induced impairment of the ubiquitin proteasome system (UPS) in SGNs

Our prior work showed that acoustic exposures at 94- or 105-dB SPL elicit pronounced increases in cochlear protein ubiquitylation accompanied by elevated levels of proteasome subunits (26). To follow up on this finding, we acquired the UPS activity reporter mouse line, Tg(CAG-Ub*G76V/GFP)1Dant, which broadly expresses the green fluorescent protein (GFP) fused to a constitutively active degradation signal (i.e., UbG76V) (32). Under normal circumstances, the GFP protein is readily ubiquitinated, marking it for degradation soon after the protein is synthesized **(Figure 4A)**. However, GFP accumulates, and fluorescence increases when the proteasome is impaired or overwhelmed. Thus, to determine which cochlear cells have impaired UPS in response to 94- or 105-dB SPL noise, we examined GFP fluorescence in cochlear mid-modiolar sections. We discovered that the GFP selectively accumulated in SGNs in a noise intensity dependent manner **(Figure 4B-C).** One potential caveat to this interpretation is that the expression of the *GFP* gene may vary among different cochlear cell types. To rule out this possibility, we examined the distribution of the *GFP* mRNA within the cochlea using RNAscope in situ hybridization. Notably, we detected *GFP* mRNA expression among various cell types within the spiral ganglion, the organ of Corti, the spiral limbus, and the inferior spiral ligament (**Figure S4**). These findings suggest that SGNs are the primarily cell type affected by proteotoxic stress in the cochlea in response to noise exposure.

**Figure 4.**
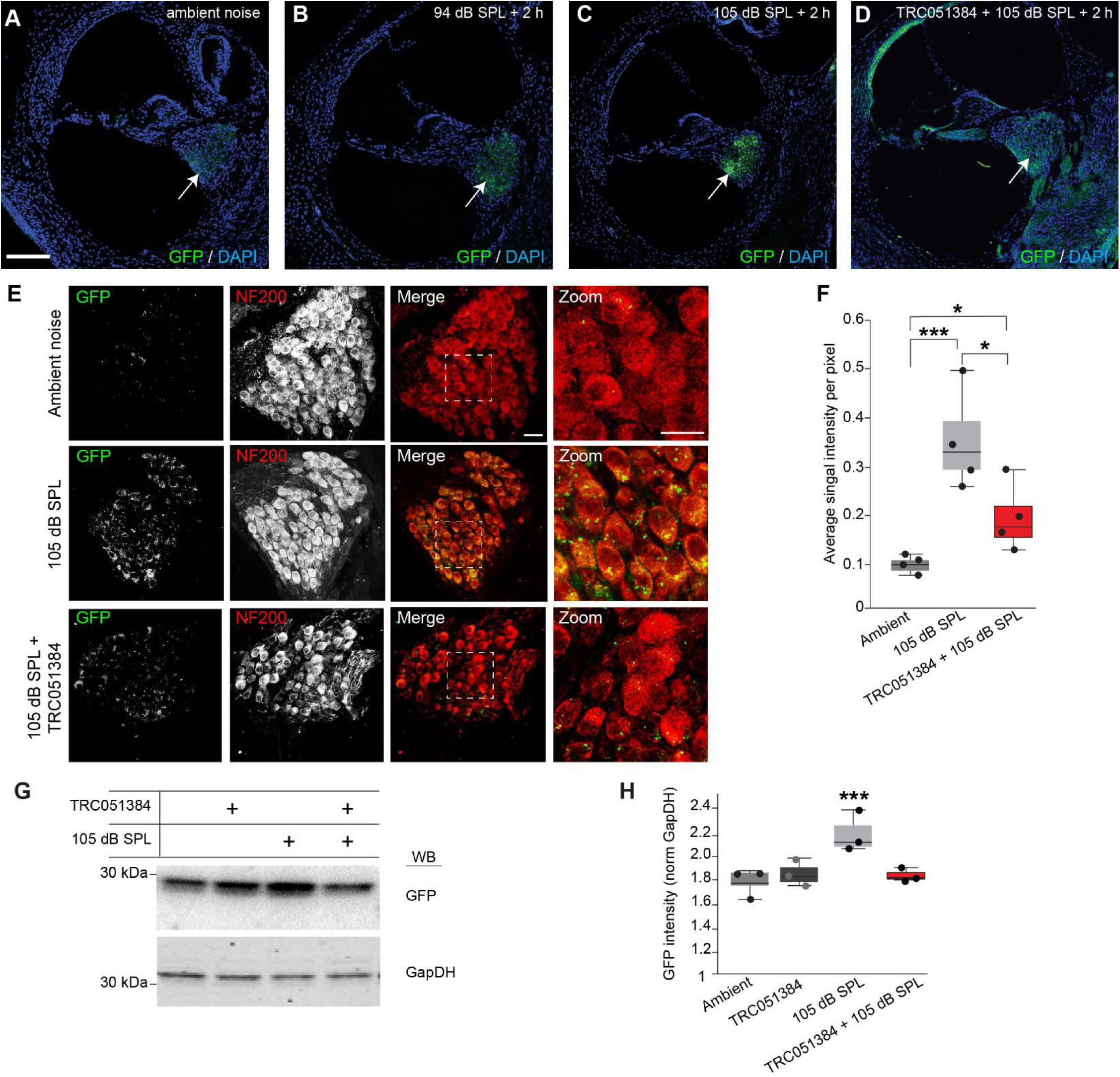
Prophylactic TRC051384 administration attenuates proteasomal impairments in C57BL/6NJ mice after 105 dB SPL noise exposure for two hours. **(A-C)** Representative fluorescent images from 12 µm cochlear sections near the mid-modiolar plane from P60 Tg(CAG-Ub*G76V/GFP)1Dant mice exposed for 2 hours to either 70 dB SPL (ambient), 94 dB SPL, or 105 dB SPL noise, followed by a 2-hour post-exposure interval. **(D)** Representative fluorescent images from 12 µm cochlear sections near the mid-modiolar plane from mice pretreated with TRC051384 and subsequently exposed to 105 dB SPL noise for 2 hours, followed by a 2-hour post-exposure interval. **(E)** Representative fluorescent images from 12 µm sections near the mid-modiolar plane, focusing on the spiral ganglion of P60 Tg(CAG-Ub*G76V/GFP)1Dant mice. As expected, noise exposure markedly increased the intensity and abundance of GFP signals within spiral ganglion neurons that were NF200 positive. Prophylactic administration of TRC051384 noticeably reduced both the size and abundance of GFP-positive clusters. **(F)** Quantification of GFP puncta intensity from (E) shows that noise exposure markedly increased puncta intensity compared to unexposed cochleae (ambient: 0.08 ± 0.024; 105 dB SPL: 0.34 ± 0.11). Prophylactic administration of TRC051384 significantly reduced GFP puncta accumulation following noise exposure, as evidenced by the lower GFP signal in the Noise + TRC051384 group (0.19 ± 0.09). **(G)** Western blot analysis of GFP protein levels in cochlear extracts from mice treated with TRC051384, with or without 105 dB SPL noise exposure for 2 hours. The upper panel shows GFP signal intensity, and the lower panel shows GAPDH as a loading control. **(H)** Quantification of (G) GFP levels normalized to the GAPDH loading control to determine relative GFP signal intensity. TRC051384 treatment alone did not significantly alter GFP abundance compared to uninjected mice (TRC051384: 1.83 ± 0.013; Uninjected: 1.77 ± 0.016). However, TRC051384 significantly reduced GFP abundance following 105 dB SPL noise exposure compared to untreated noise-exposed mice (TRC051384 + Noise: 1.82 ± 0.002; Uninjected + Noise: 2.17 ± 0.031). (F) N = 4 P60-65 mice of mixed sex per condition, (G) N = 3 P60-65 mice of mixed sex per condition. Data are represented as mean ± SD, * = p value < .05, *** = p value < .001 by one-way ANOVA determined significance with Tukey’s post hoc correction. (A-D) Scale bar = 25 µm (E) Scale bar = 15 µm.

We next examined whether prophylactic administration of TRC051384 could reduce GFP accumulation. In line with its role as a heat shock response activator, TRC051384 significantly attenuated noise-induced UPS dysfunction, as reflected by diminished GFP signal intensity in the spiral ganglion **(Figure 4D)**. To confirm that the GFP signal originated from the somata of SGNs, we performed immunostaining on the same sections using NF200 antibodies and observed strong colocalization **(Figure 4E)**. Quantitative analysis revealed that GFP signal intensity was significantly reduced in mice treated with TRC051384 before noise exposure (0.12 ± 0.048) relative to vehicle-treated controls (0.37 ± 0.10) (**Figure 4F**). Further validation of the results was achieved through western blot analysis, which showed that TRC051384 reduces the accumulation of GFP in cochlear extracts after noise exposure. Treated mice exhibited a reduced signal intensity (1.82 ± 0.002) compared to untreated controls (2.17 ± 0.03) (**Figure 4G-H**). These findings together demonstrate that the administration of TRC051384 as a prophylaxis activates the heat shock pathway, preserving the function of the UPS in the cochlea following noise exposure.

### Administration of TRC051384 prior to noise exposure protects against hearing loss

Hsf1 expression was previously found to be necessary for threshold recovery following noise exposure (33). However, the therapeutic potential of activating Hsf1 with small molecules to prevent hearing loss has received little attention. To explore this possibility, we investigated whether TRC051384-mediated activation of the Hsf1 pathway prior to noise exposure could confer protection in 16-week-old CBA/CaJ mice. TRC051384 (60 mg/kg) or vehicle alone was injected intraperitoneally 2 hours before exposure to 100 dB SPL noise for 2 hours, and ABRs were measured at 1, 3, and 14 DAN. Acute threshold shifts were observed in the TRC051384 and vehicle-treated mice at 1 DAN (**Figure 5A**). In the vehicle treated cohort at 3 DAN, thresholds remained significantly elevated at frequencies of 24 kHz and greater, and remained slightly elevated at 14 DAN. However, in the TRC051384 treated cohort threshold levels fully recovered by 3 DAN at all frequencies except 32 kHz, which then fully recovered by14 DAN. ABR wave-I amplitudes in the vehicle cohort generally failed to recover at intensities > 80 dB SPL at 16 and 32 kHz by 14 DAN **(Figure 5B-D)**. In comparison, in the TRC051384 treated cohort, ABR wave-I amplitudes fully recovered by 3 DAN. These findings were corroborated by an independent replication cohort of 16-week-old CBA/CaJ mice, which revealed that TRC051384 treatment resulted in markedly greater Wave I amplitude recovery compared to vehicle controls by 14 DAN at 24 and 32 kHz **(Figure S5A, B)**. To evaluate whether TRC051384 protects cochlear synapses, we compared ribbon synapse density between drug-treated and vehicle control groups by immunostaining cochlear whole mounts at 14 DAN with CtBP2 and GluA2 antibodies (**Figure 5E**). Consistent with TRC051384’s ability to mitigate noise-induced reductions in Wave I amplitude, the decrease in synapse density observed in vehicle-treated CBA/CaJ mice at 14 DAN was absent in TRC051384-treated mice **(Figure 5F)**. In summary, prophylactic treatment with TRC051384 preserves cochlear synapse density and auditory nerve function, supporting its potential as a protective intervention against noise-induced hearing loss.

**Figure 5.**
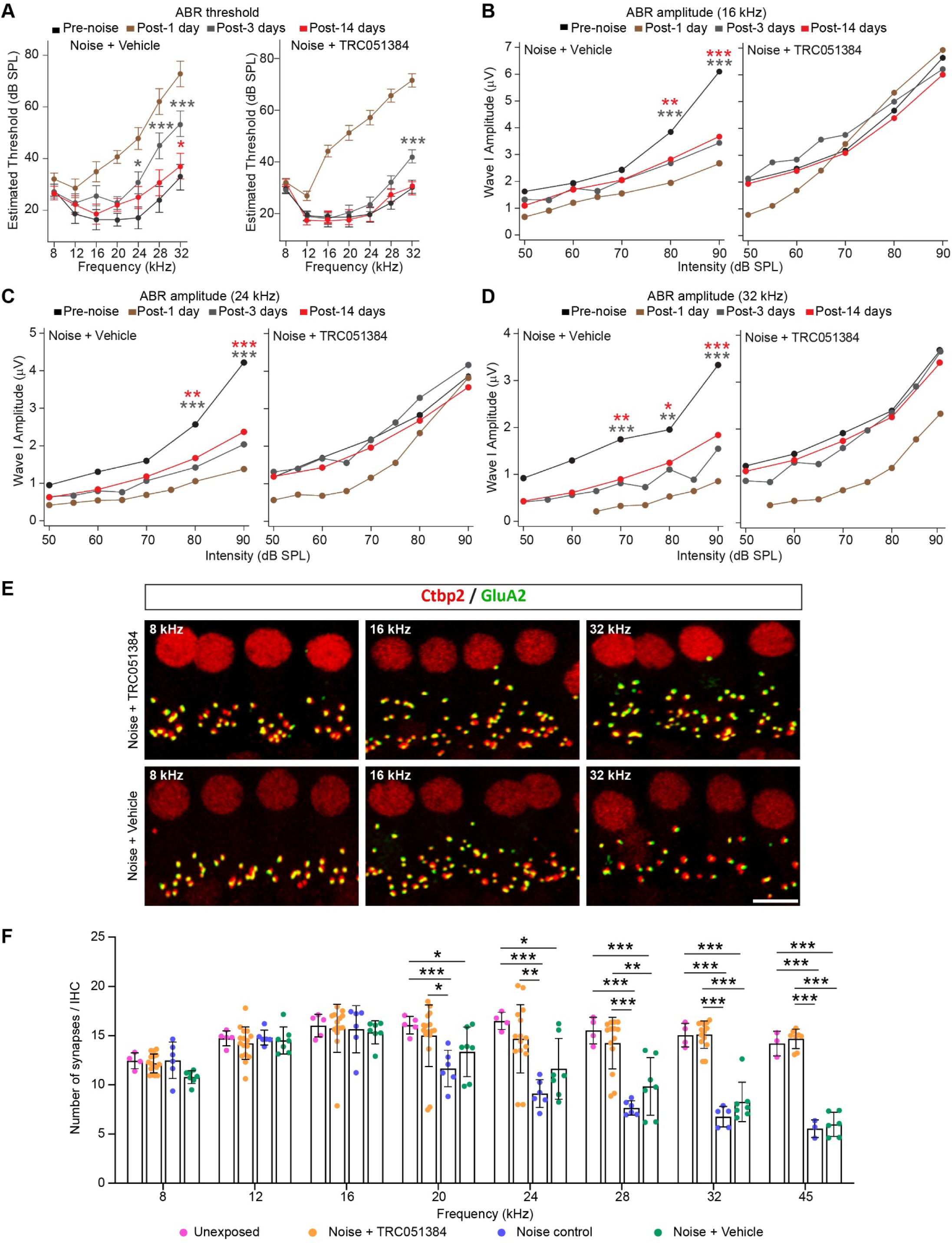
Prophylactic administration of TRC051384 accelerates recovery from noise exposure and prevents cochlear synaptopathy in male 16-week-old CBA/CaJ mice. **(A)** Auditory brainstem response (ABR) thresholds in vehicle-treated mice failed to recover to before noise (BN) levels by 3 days after noise (DAN) and did not fully recover at 14 DAN at 32 kHz (Vehicle, 14 DAN: 8 kHz = −1.42 dB; adjusted 95% CI [−8.09, 5.24], 16 kHz = 2.14 dB; adjusted 95% CI [−8.95, 13.24], 32 kHz = 4.28 dB; adjusted 95% CI [−9.86, 18.43]). In contrast, TRC051384-treated mice recovered to BN levels by 3 DAN at 8 and 16 kHz, with only partial recovery at 32 kHz (TRC051384, 3 DAN: 8 kHz = −2.35 dB; adjusted 95% CI [−6.35, 0.86], 16 kHz = 0.88 dB; adjusted 95% CI [−5.33, 7.09], 32 kHz = 11.76 dB; adjusted 95% CI [4.76, 18.77]). Complete threshold recovery was observed in TRC051384-treated mice by 14 DAN at all tested frequencies. **(B)** At 16 kHz, Wave I amplitudes in vehicle-treated mice were permanently reduced at 80- and 90-dB SPL, showing minimal recovery from 3 DAN to 14 DAN. At 3 DAN, reductions were −0.97 µV (adjusted 95% CI [−1.81, −0.14]) at <u>80 dB SPL</u> and −2.41 µV (adjusted 95% CI [−3.24, −1.58]) at <u>90 dB SPL</u>. At 14 DAN, amplitudes remained similarly reduced: −1.03 µV (adjusted 95% CI [−1.86, −0.19]) at <u>80 dB SPL</u> and −2.30 µV (adjusted 95% CI [−3.17, −1.51]) at <u>90 dB SPL</u>. In contrast, TRC051384-treated mice fully recovered Wave I amplitudes by 3 DAN. **(C)** At 24 kHz, Wave I amplitudes in vehicle-treated mice were permanently reduced at 80- and 90-dB SPL, with minimal recovery from 3- to 14-DAN. At 3 DAN, reductions were −0.92 µV (adjusted 95% CI [−1.53, −0.31]) at <u>80 dB SPL</u> and −0.69 µV (adjusted 95% CI [−2.52, −1.30]) at <u>90 dB SPL</u>. At 14 DAN, amplitudes remained reduced at −1.91 µV (adjusted 95% CI [−1.31, −0.082]) at <u>80 dB SPL</u> and −1.58 µV (adjusted 95% CI [−2.19, −0.97]) at <u>90 dB SPL</u>. In contrast, TRC051384-treated mice fully recovered Wave I amplitudes by 3 DAN. **(D)** At 32 kHz, Wave I amplitudes in vehicle-treated mice were permanently reduced at 70-, 80-, and 90-dB SPL, with minimal recovery from 3-DAN to 14-DAN. At 3 DAN, reductions were −0.82 µV (adjusted 95% CI [−1.43, −0.20]) at <u>70 dB SPL</u>, −0.75 µV (adjusted 95% CI [−1.37, −0.14]) at <u>80 dB SPL</u>, and −1.70 µV (adjusted 95% CI [−2.31, −1.08]) at <u>90 dB SPL</u>. At 14 DAN, amplitudes remained reduced at −0.76 µV (adjusted 95% CI [−1.38, −0.14]) at <u>70 dB SPL</u>, −0.69 µV (adjusted 95% CI [−1.31, −0.06]) at <u>80 dB SPL</u>, and −1.34 µV (adjusted 95% CI [−1.97, −0.73]) at <u>90 dB SPL</u>. In contrast, TRC051384-treated mice fully recovered Wave I amplitudes by 3 DAN. **(E)** Representative images from the 8, 16, and 32 kHz regions of the cochlea immunostained for CtBP2 (red) and GluA2 (green) in TRC051384- and vehicle-treated mice at 14 days after noise (DAN). Scale bar = 10 µm. **(F)** Quantification of (E) revealed significant reduction in synaptic density (based on the number of colocalized CtBP2 and GluA2 puncta) 14 DAN in vehicle-treated mice across most frequency regions of the cochlea, with the exception of the 12 and 16 kHz regions: <u>8 kHz</u>: TRC051384 12.18 ± 0.95 , Vehicle 10.78 ± 0.66; <u>20 kHz</u>: TRC051384 14.99 ± 3.12 , Vehicle 13.34 ± 2.52; <u>24 kHz</u>: TRC051384 14.68 ± 3.47, Vehicle 11.6 ± 3.08; <u>28 kHz</u>: TRC0584 14.24 ± 2.64, Vehicle 9.84 ± 2.93; <u>32 kHz</u>: TRC051384 15.09 ± 1.38, Vehicle 8.26 ± 2.01; <u>45 kHz</u>: TRC051384 14.67 ± 0.98, Vehicle 5.99 ± 1.23. (A-D) N = 17 TRC051384 treated male and 7 vehicle-treated male CBA/CaJ mice. (E-F) N = 14-17 TRC051384 treated male mice and 6-7 vehicle-treated male mice. (A-D) adjusted p value * = < .05, ** = <.01, *** = < .001; (F) Data are represented as mean ± SD.* = p value < .05, ** = p value <.01, *** = p value < .001 by one-way ANOVA with Tukey post-hoc correction.

## DISCUSSION

These findings offer new insight into previously unrecognized mechanisms of cochlear proteostasis underlying NIHL. We first confirmed that a widely used noise exposure protocol caused nearly pure TTS in 16-week-old CBA/J mice (22). Then, we performed bulk discovery-based RNA sequencing of cochlear extracts to determine the temporal dynamics of noise-induced gene expression. We detected a pronounced transcriptional response four hours after exposure, which represented the peak of gene expression. DAVID enrichment analysis identified the endoplasmic reticulum (ER) as the intracellular structure where the most significantly upregulated gene enrichment occurred (159.7-fold enrichment; *p* = 8.95 × 10⁻¹¹; FDR = 1.09 × 10⁻□), indicating that ER stress is the most prominently engaged cellular response based on gene expression changes. This enrichment was driven by elevated expression of *Hsp90b1* (Endoplasmin), *Hspa5* (BiP), *Dnajb11* (ERdj3), *Dnajc13* (RME-8), and *Sdf2l1* (BiP co-chaperone). Among these, *Hsp90b1*, *Hspa5*, *Dnajb11*, and *Sdf2l1* encode ER-resident chaperones and co-chaperones involved in protein folding and the UPR, while *Dnajc13* is implicated in endosomal trafficking and membrane dynamics. Spatial transcriptomic profiling revealed that the proteotoxic stress response was particularly pronounced in SGNs and their neighboring supportive cells. These cells exhibited elevated expression of genes encoding proteasome subunits, suggesting enhanced proteostasis activity.

To evaluate the potential protective effects of prophylactic HSR activation against noise-induced cochlear injury, we used TRC051384, a previously characterized Hsf1 activator, which was administered prior to noise exposure. Using a fluorescent UPS reporter mouse model, we mapped the prominent localization of proteostatic stress and found it to be concentrated in the spiral ganglion. Notably, TRC051384 pre-treatment reduced accumulation of the GFP-based UPS reporter, suggesting improved proteasomal function and overall protein quality control under noise-induced stress conditions. Finally, this pharmacological intervention produced functional and structural benefits. ABR Wave I amplitudes showed enhanced recovery, and synaptic integrity was better preserved. Taken together, these results underscore the potential of heat shock pathway modulation as a prophylactic strategy against NIHL.

Our findings represent a significant advance in understanding a key mechanism underlying NIHL, while also building on our and others prior work in the field. Pioneering work from over 20 years ago first identified a link between noise trauma and the proteostasis network through increased gene expression of the E3 ubiquitin ligase *Ube3b* (34). Furthermore, UBE3B is selectively expressed in SGNs in the human cochlea, supporting our finding that SGNs are a key cell type experiencing proteotoxic stress following noise exposure (35). Further evidence supporting the involvement of noise-induced impaired proteostasis, specifically in SGNs, comes from single-cell transcriptomic analyses. These analyses identified *DDIT3* (i.e., *CHOP*), a key transcription factor that regulates ER stress and apoptosis, as being significantly elevated in this cell population (36). Our discovery that activating the HSR with TRC051384 mitigates noise-induced damage aligns with earlier findings that report acutely elevating body temperature in CBA/CaJ mice significantly reduces noise-induced threshold shifts via HSR activation (37). Our findings also support and extend previous work demonstrating the importance of ER stress and the activation of the UPR in response to noise exposure, particularly through the ER-resident protein Tmtc4 (38). Furthermore, prior studies have shown that pharmacological activation of eIF2B, which suppresses the UPR, can mitigate NIHL (38, 39). More recently, mutations in the transcription factor ATF6, which regulates a gene expression program critical for maintaining ER function and initiating the UPR, have been linked to a deafness-blindness syndrome (40, 41).

As with any study, our work has limitations that warrant consideration. First, the initial RNA sequencing analysis revealed strong, specific upregulation of genes encoding components of the ER chaperone complex. However, this bulk approach is biased toward changes in gene expression in the most abundant cochlear cell types, which limits resolution. A key limitation of our spatial transcriptomic analysis was the absence of probes targeting many ER chaperone genes. This prevented us from directly assessing their expression at the level of individual cell types. Despite considerable effort, we were unable to obtain sufficient transcriptomic signal from IHCs, OHCs, and supporting cells within the organ of Corti, likely due to the platform’s limited sensitivity (42). However, in our previous studies, we showed that both Hsp90b1 and Psmc5 have elevated levels in hair cells based on immunofluorescent microscopy (26). It is also important to note that while our previous findings showed a noise intensity–dependent increase in the abundance of many proteins (26), suggestive of overproduction or stress-induced misfolding, additional cellular stressors likely contribute to this response. In particular, oxidative stress and disrupted calcium homeostasis are well-known inducers of ER chaperone expression and likely contribute in shaping the complex, multifactorial response to auditory trauma (43). While we demonstrated that TRC051384 activates expression of the Hsf1 target gene, *Hspa1a,* in both heterologous cells and cochlear organotypic cultures, we acknowledge that we did not directly confirm activation of the HSR using qPCR *in vivo* within the intact mouse cochlea. Although the UPS fluorescent reporter mouse offered a sensitive tool to identify cells experiencing proteotoxic stress and to assess the mitigating effects of TRC051384, it is important to acknowledge that this represents an artificial reporter system.

Looking ahead, several key avenues should be pursued to build on our findings. First, although our results revealed that SGNs are a key cell type undergoing proteotoxic stress following acoustic trauma, further studies are needed to confirm and characterize this response across additional cochlear cell types. Second, additional mechanistic studies are needed to understand how the heat shock response intersects with other cellular stress pathways such as the UPR, and how it integrates stress responses, oxidative stress, calcium dysregulation, and mitochondrial dysfunction. Third, although TRC051384 showed promising protective effects in mice, future studies will be needed to evaluate their pharmacokinetics, cochlear bioavailability, and potential off-target effects. Testing this compound or improved Hsf1 modulators in other mouse strains and in larger animal models (e.g., guinea pigs or non-human primates) is a critical next step for advancing its translational potential. Additionally, studies exploring optimal timing, dosage, and delivery routes are also necessary to maximize therapeutic efficacy and minimize risk. Finally, future studies should investigate whether activating the HSR after noise exposure can confer therapeutic benefits. These efforts will collectively help to clarify the feasibility of targeting cochlear proteostasis as a preventive or therapeutic strategy for noise-induced hearing loss in humans.

## METHODS

### Sex as a biological variable

Mice and rats of mixed sexes were used in our experiments; however, the presented results were not sufficiently powered to determine sex-specific effects. Male mice were used in Figures 1 and 5.

### Mouse procedures

Auditory brainstem responses (ABRs) and distortion product otoacoustic emissions (DPOAEs) were recorded from young adult mice (postnatal day ∼56 or ∼112). All recordings were performed in a sound-attenuated chamber (Eckel Audiometric Room, Cambridge, MA) using a Tucker-Davis Technologies (TDT) System III workstation running BioSigRP software. ABR stimuli were pre-amplified using a 4-channel MEDUSA preamplifier and processed in real-time by the TDT System III. Prior to electrode placement, animals were anesthetized via intraperitoneal injection of ketamine (100 mg/kg) and xylazine (3 mg/kg). Supplemental injections (25% of the initial dose) were administered as needed to maintain anesthesia. Core body temperature was maintained between 37–38°C using an electric heating pad (Homeothermic Blanket System, Harvard Apparatus). Platinum subdermal needle electrodes were placed at the vertex (recording), ipsilateral mastoid (reference), and lower back (ground). ABRs were evoked by 20-millisecond tone pips (8–32 kHz, 4 kHz increments) at sound levels from 10 to 90 dB SPL, with 5 dB steps from 10–40 dB and 10 dB steps from 40–90 dB. Responses were averaged across 500 repetitions. ABR thresholds were defined as the lowest sound level eliciting a detectable Wave I with an amplitude at least 0.18 μV above the noise floor. Wave I amplitude was measured as the voltage difference between the Wave I peak (P1) and the following trough (N1). For DPOAE recordings, an Etymotic ER-10B+ low-noise microphone system was used. Stimuli consisted of two pure tones (f1 and f2), differing by a factor of 1.2. L1 and L2 levels were swept from 15 to 75 dB SPL (with L1 > L2 by 5 dB) in 5 dB increments across frequencies from 8 to 32 kHz. The spectral magnitudes of f1, f2, the 2f1–f2 distortion product, and the noise floor were derived by averaging the responses to 500 presentations.

Noise exposures were delivered to unanesthetized animals placed in small wire cages within a custom-built sound-attenuated chamber designed by Charles Liberman (Mass. Eye and Ear). The chamber was constructed from 3/4" plywood sheets, with the key design principle being that no two sides were parallel, minimizing acoustic reflections. The front and back panels were identical in shape, except that the rear panel was truncated at a height of 42", creating a slanted top surface relative to the floor. Side panels were cut to fit this asymmetrical geometry. The top panel included a rectangular cutout for mounting an exponential horn and acoustic driver. Acoustic overexposure waveforms were generated using a Tucker-Davis Technologies waveform generator (Alachua, FL), producing band-limited noise spanning 8–16 kHz for a total duration of 30 minutes or two hours. Noise was delivered via an exponential horn connected to a JBL 2446H/J compression driver (Northridge, CA). Mice were exposed to calibrated noise levels ranging between 94- and 105-dB SPL (1.00 Pa Sound Intensity Level, SIL). Sound pressure level stability was monitored in real time using a PicoLog system (Picoscope 2000 series) and a calibrated microphone (PCB Piezotronics, NY).

B6.Cg-Tg(CAG-Ub*G76V/GFP)1Dant/J, (Strain #:008111, RRID:IMSR_JAX:008111) were purchased from Jax.

TRC051384 was prepared in a vehicle consisting of 12% dimethyl sulfoxide (DMSO) and 88% vehicle #1 (10% DMSO/30% Kolliphor EL/10% Ethanol/50% PBS) and the solution was filtered to sterilize. The compound was administered to mice via intraperitoneal injection (IP) at a dose of 60 mg/kg. Injections were carried out under aseptic conditions using standard IP techniques to ensure accuracy and reproducibility.

### Bulk RNA-Sequencing and qPCR

For Bulk RNA-sequencing, total RNA was extracted from cochlear tissue using TRIzol reagent (Thermo Fisher) according to the manufacturer’s protocol and further purified with the RNeasy Plus Mini Kit (Qiagen). Purified RNA was sent to Novogene (Sacramento, CA, USA) for quality control (QC), library preparation, sequencing, and quantitation according to their standard gene expression workflow. RNA degradation and contamination were initially assessed on 1% agarose gels. RNA purity was evaluated using a NanoPhotometer spectrophotometer (IMPLEN, CA, USA), while RNA integrity and concentration were determined using the RNA Nano 6000 Assay Kit with the Bioanalyzer 2100 system (Agilent Technologies, CA, USA). Only RNA samples that passed QC were used for library preparation and sequencing.

For library construction, 1 µg of total RNA per sample was used as input. Libraries were prepared with the NEBNext Ultra™ RNA Library Prep Kit for Illumina (NEB, USA) according to the manufacturer’s instructions, with unique index codes added to each sample. Briefly, mRNA was enriched using poly-T oligo-attached magnetic beads, fragmented under elevated temperature in First Strand Synthesis Reaction Buffer (5X), and reverse-transcribed using random hexamer primers and M-MuLV Reverse Transcriptase (RNase H–). Second-strand cDNA synthesis was performed with DNA Polymerase I and RNase H, followed by end-repair and adaptor ligation with NEBNext adaptors containing a hairpin loop structure. cDNA fragments of 150–200 bp were size-selected using the AMPure XP system (Beckman Coulter, USA). USER Enzyme (NEB, USA) treatment was performed at 37 °C for 15 min, followed by 95 °C for 5 min. Libraries were amplified with Phusion High-Fidelity DNA Polymerase, Universal PCR primers, and Index primers. PCR products were purified with AMPure XP, and library quality was verified on the Agilent Bioanalyzer 2100 system. Indexed libraries were clustered on a cBot Cluster Generation System with PE Cluster Kit cBot-HS (Illumina) and paired-end sequencing (2 × 150 bp) was performed on an Illumina NovaSeq 6000 platform.

Raw FASTQ reads were processed with fastp to remove adaptor sequences, poly-N reads, and low-quality reads. Clean data quality metrics (Q20, Q30, and GC content) were assessed. Reference genome and annotation files were obtained from NCBI/UCSC/Ensembl. Genome indexing and alignment were performed with STAR (v2.5) using the Maximal Mappable Prefix (MMP) algorithm, achieving >95.3% total mapping efficiency and >75 million mapped reads per sample across four biological replicates per group. Gene-level counts were obtained with HTSeq (v0.6.1).

Differential gene expression analysis was conducted using iGEAK (R/Shiny-based pipeline). Low-abundance transcripts (read count < 8 in fewer than 3 samples) were filtered out. Data were normalized using edgeR’s TMM (Trimmed Mean of M-values) method. Statistical testing for differential expression was performed with edgeR, and p-values were adjusted for multiple testing with the Benjamini–Hochberg method to control the false discovery rate (FDR). Genes with adjusted p-value < 0.05 and fold-change > 1.5 were considered significantly differentially expressed.

Quantitative PCR (qPCR) was performed as follows. Total RNA was isolated using TRIzol reagent (Thermo Fisher, Cat. #15596026) and purified with the RNeasy Kit (Qiagen, Cat. #74104). cDNA was synthesized from isolated RNA using the iScript cDNA Synthesis Kit (Bio-Rad, Cat. #1708890). qPCR was carried out on a QuantStudio 3 Real-Time PCR System (Thermo Fisher, Cat. #A28567) using Power SYBR Green Master Mix (Thermo Fisher, Cat. #A46109). Each reaction contained 50 ng of cDNA. The thermal cycling conditions were as follows: initial denaturation at 95 °C for 10 min, followed by 40 cycles of 95 °C for 15 s and 60 °C for 60 s. Primers were purchased from IDT and reconstituted to a 10 µM working concentration. The following primer sequences were used: Hsf1 forward, ATGCCATGGACTCCAACCTG; Hsf1 reverse, CTCCTGAATGTCCAGCAGGG; Hspa1a forward, TGATCGCCGGTCTAAACGTG; Hspa1a reverse, CCCAGGTCGAAGATGAGCAC.

### Spatial transcriptomics

Spatial transcriptomic profiling was conducted using the NanoString GeoMx Digital Spatial Profiler at the Northwestern University Immunotherapy Assessment Core (IAC) following the manufacturer’s guidelines. Mouse cochleae were recovered from sham or treated mice and embedded in OCT. Cochleae were sectioned at a thickness of 6 μm and processed for the GeoMx Mouse Whole Transcriptome Atlas (cat# GeoMx NGS RNA WTA Mm, Nanostring, Bruker spatial biology) according to the manufacturer’s instructions with an exception of 0.0001% Proteinase K used in slide preparation step. Regions of interest (ROI) were selected in the “tissue area name”. Cells were collected in this order: (1) NF200+, and (2) NF200- cells with a target of at least 50 events per ROI. Segmentation was performed with the following settings: 0 Erode, 2 µm N-Dilate, 100 µm2 Hole Size, and 5 µm2 Particle Size. After construction and quality control (Qubit and bioanalyzer) of the GeoMx library, single-end 50 nucleotide sequencing was performed on a Hiseq 4000 System (Illumina) with a total of 100 million reads. After sequencing, the NanoString GeoMx Digital Spatial Profiler (DSP) with DRAGEN pipeline were used to automatically process the Illumina FASTQ sequencing files to GeoMx readable digital counts conversion files for input back into the DSP Data Analysis Suite. Data comparisons represent total cells pooled from 21 ROIs for the noise exposed group and 30 ROIs from the control group.

### NGS library preparation and sequencing

Quality control for the constructed library was performed using the Agilent Bioanalyzer High Sensitivity DNA kit (Agilent Technologies, 5067-4626) and the Qubit DNA HS assay kit for qualitative and quantitative analysis, respectively. A Qubit concentration of 0.156 ng/µL had an average fragment size of 182 bp. The final library was sequenced on Illumina HiSeq4000 sequencer using single end 50nt mode.

### Mass spectrometry analysis

Cochlear extracts were prepared as follows. Two cochleae from control 4-month-old C57BL/6NJ mice were solubilized in 100 µL of 2% acetonitrile (ACN) buffer containing 0.05% trifluoroacetic acid (TFA) using a Precellys homogenizer and incubated on ice for five minutes. Samples were centrifuged at 12,000 × g for 15 min, and the supernatant transferred to a fresh tube. Four volumes of ACN were added, mixed thoroughly, and incubated on ice for 5 min. Following a second centrifugation at 12,000 × g for 15 min, the supernatant was collected and dried in a SpeedVac concentrator. The dried pellet was resolubilized in 50 µL of buffer A (94.785% H_₂_O, 5% ACN, 0.125% formic acid [FA]), and 15 µL was injected for LC–MS/MS analysis. Samples were loaded via autosampler using a Thermo EASY-nLC 1000 UPLC pump onto a vented Acclaim PepMap 100 trap column (75 µm × 2 cm, Thermo Fisher Scientific) connected to an Acclaim PepMap 100 nanoViper analytical column (75 µm × 500 mm, 3 µm particle size, 100 Å pore size, C18; Thermo Fisher Scientific, Cat# 164570) with a stainless-steel emitter tip. The column was interfaced with a Nanospray Flex Ion Source operating at a spray voltage of 2000 V. Mass spectra were acquired on an Orbitrap Fusion mass spectrometer (Thermo Fisher Scientific). Buffer A consisted of 94.785% H_₂_O, 5% ACN, and 0.125% FA, while buffer B consisted of 99.875% ACN with 0.125% FA. Peptides were separated over a 60-min gradient with the following profile: 2–8% buffer B (3 min), 8–24% (32 min), 24–36% (10 min), 36–55% (5 min), 55–95% (5 min), followed by 95% (5 min).

We analyzed 50 pg of TRC051384 (TRC051384: CAS No.: 867164-40-7) resuspended in Buffer A. MS methods were as follows. Targeted SIM (tSIM) was used for quantification. For MS1, quantitation was performed on the precursor ion at *m/z* 465.23 (233.6261, +2). For MS2, quantitation was performed on the same precursor with multiple fragment ions analyzed. MS data were manually inspected using Excalibur software, and MS2 spectra were further processed with Compound Discoverer (Thermo Scientific). For MS1, precursor ions were isolated using the quadrupole with a 20 *m/z* isolation window, and detection was performed in the Orbitrap at 120,000 resolution (positive ion mode). For MS2, precursor isolation was carried out with a 1.6 *m/z* isolation window, followed by HCD fragmentation at 30% normalized collision energy. Detection was performed in the Orbitrap at 30,000 resolution with RF Lens set to 30% (positive ion mode).

### Western Blots

Cochleae were rinsed in PBS and homogenized in Syn-PER buffer (Thermo Scientific, Cat# 87793) supplemented with 1% Triton X-100 (Sigma-Aldrich, Cat# T8787) and 0.5% SDS using a Percellys 24 tissue homogenizer (Bertin, Cat# P000669-PR240-A). Homogenization was performed at room temperature with three 30-second pulses at 6800 rpm. Lysates were incubated on ice for 10 min. Samples were then incubated on ice for an additional 30 min, followed by centrifugation at 10,000 × g for 5 min at 4 °C. The resulting supernatants were collected, and protein concentrations were determined using the Pierce BCA Protein Assay Kit (Thermo Scientific, Cat# 23225). For immunoblotting, 25 µg of cochlear lysate was mixed with 6× loading buffer, boiled at 100 °C for 5 min, and immediately loaded onto a 10% Tris gel (Bio-Rad). Proteins were electrophoretically separated and transferred to nitrocellulose membranes using the Bio-Rad Trans-Blot Turbo transfer system (1.3 A, 25 V, 10 min). Membranes were blocked with 1× Odyssey blocking buffer (LI-COR, Cat# 927-70001) for 1 h at room temperature, then incubated overnight at 4 °C with primary antibodies GFP (1:500, Sigma, Cat#: SAB4301138, RRID: AB_2750576) and GAPDH (Cell Signaling Technology, Cat#: 2118S, RRID: AB_561053). After three washes in TBST, membranes were incubated with secondary antibodies for 2 h at room temperature. Blots were imaged using the Odyssey DLx Imaging System (LI-COR).

### Fluorescent microscopy

Mice were sacrificed, and temporal bones / cochleae were isolated within 3 minutes of euthanasia. Samples were rinsed in ice-cold PBS supplemented with Halt protease inhibitor cocktail until the buffer remained clear of blood after vortexing. The oval and round windows were punctured, and a small hole was made near the apex of the cochlea to gently perfuse the scalae with PKS solution (4% paraformaldehyde, 126 mM NaCl, 2.5 mM KCl, 25 mM NaHCO_₃_, 1.2 mM NaH_₂_PO_₄_, 1.2 mM MgCl_₂_, 2.5 mM CaCl_₂_, 11 mM sucrose). Samples were then immersed in PKS and fixed for 1 h at 4 °C. Cochleae were decalcified in Immunocal (Decal Chemical Corp., Congers, NY) for ∼2 h at 4 °C, or until translucent. The decalcified bone, lateral wall, Reissner’s membrane, and tectorial membrane were carefully dissected away from the organ of Corti. For whole-mount preparations, cochleae were divided into four pieces, cryoprotected in 30% sucrose for 30 min, and frozen on dry ice. Frozen tissues were thawed and rinsed three times in PBS for 10 min each at room temperature with agitation.

Cochlear pieces were blocked overnight at 4 °C in blocking buffer (20% normal horse serum, 1% Triton X-100 in PBS). Primary antibodies were diluted in blocking buffer as follows: Recombinant Neurofilament H (NF200) (1:1000, Synaptic Systems, Cat# 171106, RRID: AB_2721078), CtBP2 (1:200, Thermo, Cat #: 612044, RRID: AB_399431), GluA2 (1:100, Chemicon, Cat #: MAB397, RRID: AB_2113875), Myo7a (1:500, Thermo, Cat #: PA1-936, RRID: AB_2235704). Samples were incubated with primary antibodies at 37 °C overnight, rinsed three times in PBS (15 min each), and then incubated with secondary antibodies for 3 h at room temperature: Goat Anti-Mouse IgG2a Alexa Fluor 488 (1:500, Abcam A21131), Goat Anti-Mouse IgG1 Alexa Fluor 568 (1:500, Abcam A21124), Goat Anti-Rabbit H&L Alexa Fluor 647 (1:250, Abcam A27040), Donkey Anti-Sheep H&L Alexa Fluor 647 (1:250, Abcam ab150179), and Donkey Anti-Sheep H&L Alexa Fluor 405 (1:250, Abcam ab175676). After secondary incubation, samples were washed three times in PBS (15 min each) and counterstained with DAPI solution (Abcam ab228549) for 5 min. Tissues were mounted using ProLong Gold Antifade Mountant (Thermo Scientific, Cat# P10144). Images were captured on a Leica DMI4000 confocal laser scanning microscope under identical acquisition settings.

### RNAscope

Mouse cochleae were harvested and dissected as described above. Cochleae were fixed for 1 h, then decalcified overnight in Immunocal. Samples were subsequently cryoprotected overnight in 30% sucrose buffer. Once saturated, cochleae were embedded in OCT and frozen at –80 °C. Tissue was sectioned at 12 µm and mounted onto frosted glass slides, which were stored at –80 °C until use.

Slides were washed three times in PBS at room temperature, followed by sequential dehydration in 30%, 50%, 70%, and 100% ethanol. Slides were then incubated with 3% hydrogen peroxide at room temperature for 10 min, treated with RNAscope Protease III for 15 min at 40 °C, and rinsed three times in DEPC-treated water. RNAscope probes were prewarmed at 40 °C in a water bath for 10 min to dissolve any precipitate, then cooled to room temperature. Target probes for C1, C2, and C3 were mixed in a 50:1:1 ratio, with a final volume of 50 µL per sample. Slides were incubated with the probe mixture for 2 h at 40 °C in the HybEZ Oven. Each channel was then amplified and hybridized to the appropriate fluorophore following the manufacturer’s instructions. Nuclei were counterstained with DAPI. Images were acquired using a confocal laser scanning microscope (Leica DMI4000) under identical acquisition settings. The mouse probes used were as follows: Hspa5 (ACD Bio-Techne, Cat#: 438831-C), Hsp90b1 ACD Bio-Techne, Cat#: 556051), Sdf2l1 (ACD Bio-Techne, Cat#: 562401), Tubb3-C2 (ACD Bio-Techne, Cat#: 423391-C2), Myo7a (ACD Bio-Techne, Cat#: 462771), and GFP-C2 - Synthetic construct Cox8ND6gfp fusion protein gene complete cds custom sequence 709/714 (99%).

### Statistics

Statistical analyses were performed using Excel, GraphPad prism, and SAS 9.4. Data are presented as mean ± SD. For measures obtained across multiple stimulus intensities or frequencies (e.g., ABR thresholds, DPOAE thresholds), comparisons were performed using one-way ANOVA. Post hoc pairwise comparisons between genotypes were corrected for multiple testing using Tukey’s method. Statistical significance was defined as an adjusted p-value < 0.05.

For the ABR analysis, we use linear mixed effects regression models (LMMs) to account for the correlation between repeated measurements within mouse over time and at different frequencies and intensities. We fit separate models for Noise + Vehicle and Noise + TRC051384. To model changes in threshold by frequency and DAN, we estimated LMMs with threshold as the outcome, fixed effects for frequency and DAN, and their interaction, and a random intercept for mouse to account for repeated measures over time. The resulting beta coefficients informed our understanding of how treatment altered hearing recovery at different frequencies over time. For changes in Wave I Amplitude by intensity and DAN, we similarly estimated LMMs with amplitude as the outcome, fixed effects for intensity and DAN, and their interaction, and a random intercept for mouse. We fit these models separately for each frequency (16, 24 and 32 kHz). Statistical comparisons in Figures 1A, 1C, 5A-D, S5A-B are all model-based.

### Study approval

All animal experiments were conducted according to Northwestern University Institutional Animal Care and Use Committee (IACUC approved protocol numbers IS00011904, IS00001182, and IS00022569). For euthanasia, pups were immediately decapitated, and adult animals were euthanized by isoflurane overdose and decapitation.

### Data availability

The RNA sequencing datasets supporting the conclusions of this study have been deposited in the NCBI Gene Expression Omnibus (GEO). Bulk RNA sequencing data are available under accession number **GSE312476**, and spatial transcriptomic data are available under accession number **GSE312646**.

## AUTHOR CONTRIBUTIONS

J.Y., M.A.R., M.A.R., M.A.C. and J.N.S., designed experiments. J.Y., and M.A.R. performed ABR and DPOAE experiments. J.Y., S.E., and M.A.R. performed bulk RNA-Seq experiments. M.A.R. performed spatial RNA-Seq experiments. S.E. performed MS analysis. M.A.R. performed the in vitro experiments.

J.Y. and M.A.R. performed the IF. M.A.R. performed the WBs. J.Y., M.A.R., Y.Z.W., S.E., M.S., L.J.W., and J.N.S. analyzed the data. M.A.R., J.Y. and J.N.S. wrote the manuscript and all authors reviewed and edited the manuscript.

## ACKNOWLEDGEMENTS

Auditory research in the Savas laboratory was supported by grants W81XWH-22-1-0773 and W81XWH-19-1-0627, and by NIDCD grant R01DC014712 awarded to M.A.R. The authors thank Ms. Chen (“Jen”) Yeh for her contributions to data collection and early versions of the ABR analyses.

