## Supplementary material for "Heat-shock pathway activation by TRC051384 protects spiral ganglion neurons from noise-induced hearing loss": Suplementary Materials

**Inventory of Supplemental Information**

Supplemental Data: 5 Supplemental Figures and Legends

Supplemental Table Legends

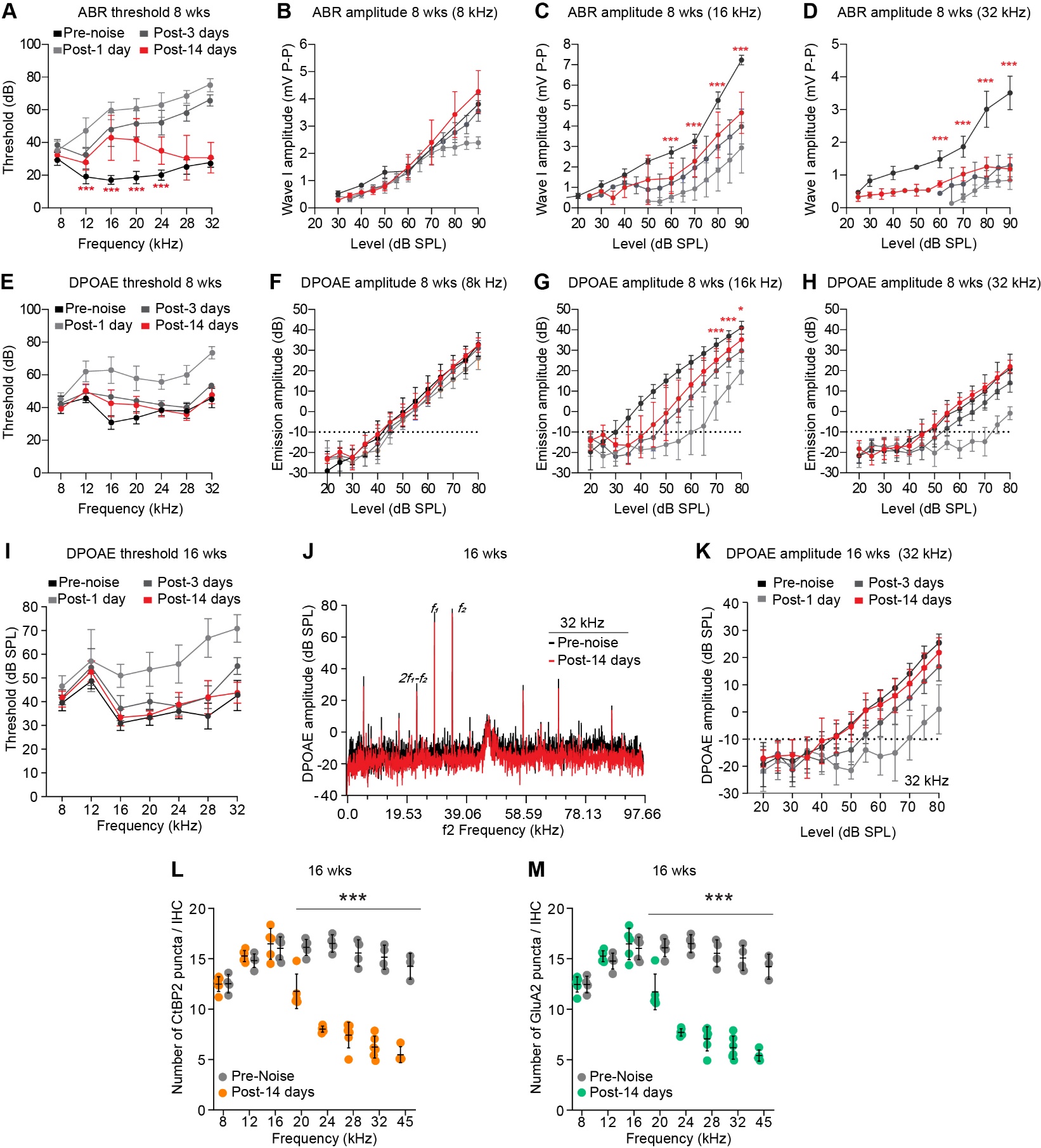

**Supplementary Figure 1. Noise exposure causes persistent auditory dysfunction and synaptic loss in 8- week-old CBA/CaJ mice 14 days after noise (DAN).**

**(A)** ABR thresholds remained significantly elevated 14 DAN compared to BN levels from 12-24 kHz in 8 week old CBA/CaJ mice (**12 kHz:** BN =17.14 ± 2.67, 14 DAN = 27.5 ± 5.24; **16 kHz:** BN = 17.14 ± 2.67, 14 DAN = 40 ± 12.65; **20 kHz:** BN = 18.57 ± 3.78, 14 DAN = 38 ± 10.37; **24 kHz:** BN = 20 ± 3.16, 14 DAN = 34 ± 8.94) N = 7. Data are represented as mean ± SD, *** = p value < .001 by one-way ANOVA determined significance with Tukey’s post hoc correction.

**(B)** Wave I amplitude at 8 kHz fully recovered to baseline levels by 14 DAN in 8-week-old male CBA/CaJ mice (N = 7).

**(C)** Wave I amplitude failed to recover to BN levels by 14 DAN at 16 kHz at intensities above 60 dB SPL (**60 dB SPL:** BN = 2.71 ± 0.28, 14 DAN = 1.45 ± 0.74; **70 dB SPL:** BN = 3.26 ± 0.34, 2.28 ± 0.76; **80 dB SPL:** BN = 5.26 ± 0.41, 14 DAN = 3.57 ± 1.11; **90 dB SPL:** BN = 7.23 ± 0.24, 14 DAN = 4.65 ± 1.01). N = 7. Data are represented as mean ± SD, *** = p value < .001 by one-way ANOVA determined significance with Tukey’s post hoc correction.

**(D)** Wave I amplitude failed to recover to BN levels by 14 DAN at 32 kHz at intensities above 60 dB SPL (**60 dB SPL:** BN = 1.48 ± 0.26, 14 DAN = 0.73 ± 0.10; **70 dB SPL:** BN = 1.86 ± 0.32, 1.03 ± 0.21; **80 dB SPL:** BN = 3.01 ± 0.55, 14 DAN = 1.25 ± 0.29; **90 dB SPL:** BN = 3.51 ± 0.51, 14 DAN = 1.18 ± 0.35). N = 7. Data are represented as mean ± SD, *** = p value < .001 by one-way ANOVA determined significance with Tukey’s post hoc correction.

**(E)** DPOAEs from the 8-32 kHz regions fully recovered to BN threshold levels by 14 DAN. N = 12 BN, 8-week-old mice.

**(F)** DPOAE amplitudes fully recovered by 14 DAN to BN levels at 8 kHz.

**(G)** DPOAE amplitudes at 16 kHz failed to recover by 14 DAN relative to BN levels at intensities above 60 dB SPL (**60 dB SPL:** BN = 24.11 ± 3.33, 14 DAN = 13.30 ± 7.49; **70 dB SPL:** BN = 32.7 ± 3.16, 25.02 ± 5.82; **80 dB SPL:** BN = 41.03 ± 3.14, 14 DAN = 35.17 ± 5.93). N = 12 BN, 7 14 DAN 8-week-old mice. Data are represented as mean ± SD, *** = p value < .001 by one-way ANOVA determined significance with Tukey’s post hoc correction.

**(H)** DPOAE amplitudes fully recovered by 14 DAN to BN levels at 32 kHz.

**(I-K)** DPOAE thresholds for f2=8-32 kHz showed a complete recovery by 14 DAN compared to BN levels. Representative DPOAE spectra and amplitude functions for f2=32 kHz highlight the full recovery of DPOAE amplitude 14 DAN. N = 6 16-week-old male CBA/CaJ mice. Adjusted p-value = > .001 data are represented as mean ± SD. *** = p value < .001 by one-way ANOVA with Tukey post hoc correction.

**(L-M)** Quantification of presynaptic ribbons (CtBP2, apricot) and postsynaptic AMPARs (GluA2, green) revealed significant reductions in the average density of both synaptic elements across the 20–45 kHz regions of the cochlea following noise exposure (CtBP2 puncta per IHC: 14 DAN = 7.78 ± 2.44, Control = 15.53 ± 0.88; GluA2 puncta per IHC: 14 DAN = 7.63 ± 2.44, Control = 15.49 ± 0.88). N = 4 unexposed, 6 exposed cochleae from individual 16-week-old male CBA/CaJ mice. *p < .001 by one-way ANOVA with Tukey’s post hoc correction.

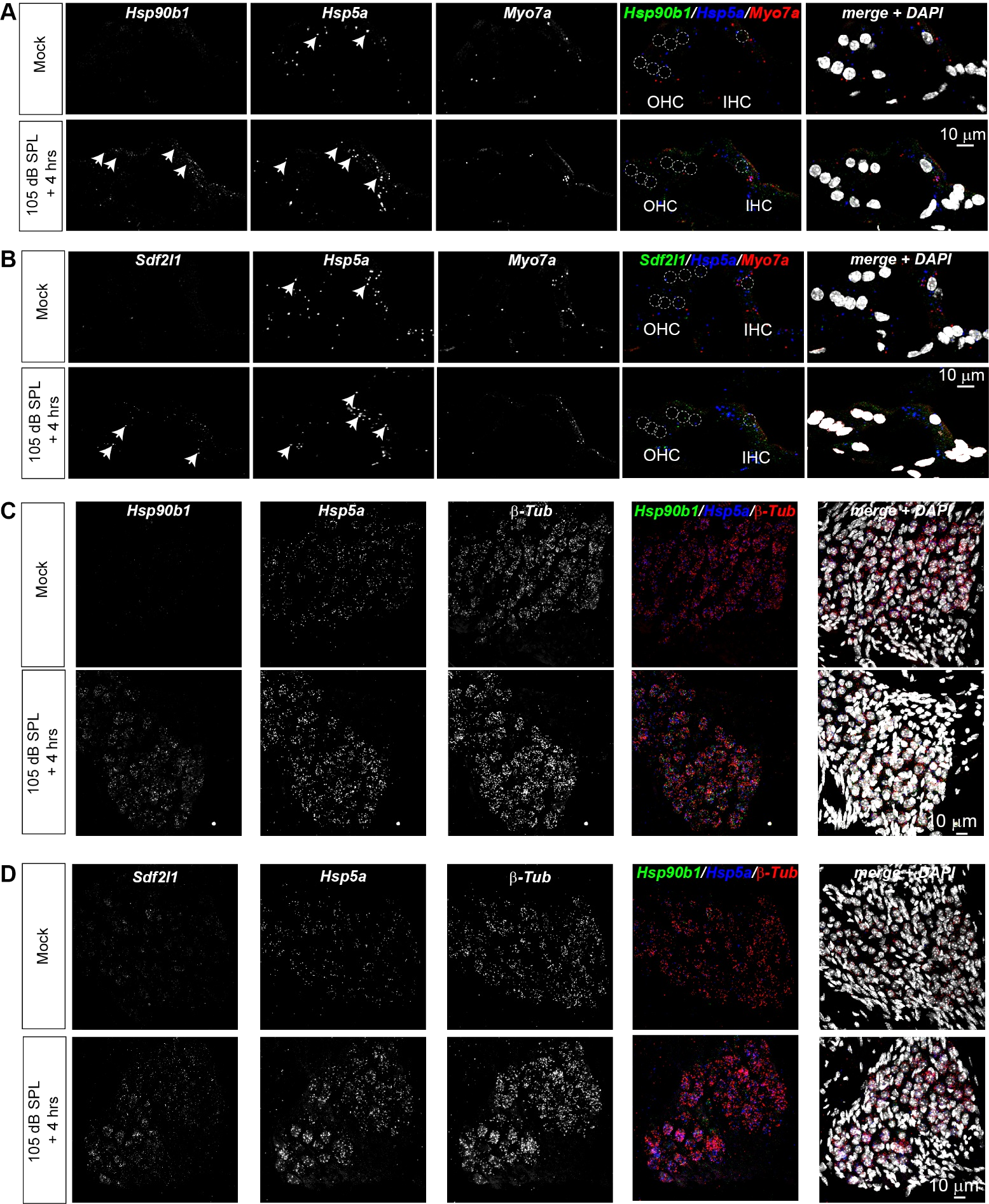

**Supplementary Figure 2. Noise exposure markedly elevated *Hsp90b1*, *Hspa5*, and *Sdf2l1* expression in the spiral ganglion and in both inner and outer hair cells in 16-week-old male CBA/CaJ mice.**

**(A)** Representative RNAscope *in situ* hybridization images of the organ of Corti showing *Hsp90b1* (green), *Hspa5* (blue), and the hair cell marker *Myo7a* (red), with signal intensity generally increased after noise exposure in and near both inner and outer hair cells.

**(B)** Representative RNAscope *in situ* hybridization images of the organ of Corti showing *Sdf2l1* (green), *Hspa5* (blue), and the hair cell marker *Myo7a* (red), with signal intensity generally increased after noise exposure in and near both inner and outer hair cells.

**(C)** Representative RNAscope *in situ* hybridization images of the spiral ganglion showing *Hsp90b1* (green), *Hspa5* (blue), and the SGN marker *β-Tub* (red), with signal intensity generally increased after noise exposure in and near SGNs.

**(D)** Representative RNAscope *in situ* hybridization images of the spiral ganglion showing *Hsp90b1* (green), *Hspa5* (blue), and the SGN marker *β-Tub* (red), with signal intensity generally increased after noise exposure in and near SGNs.

Scale bars = 10 μm.

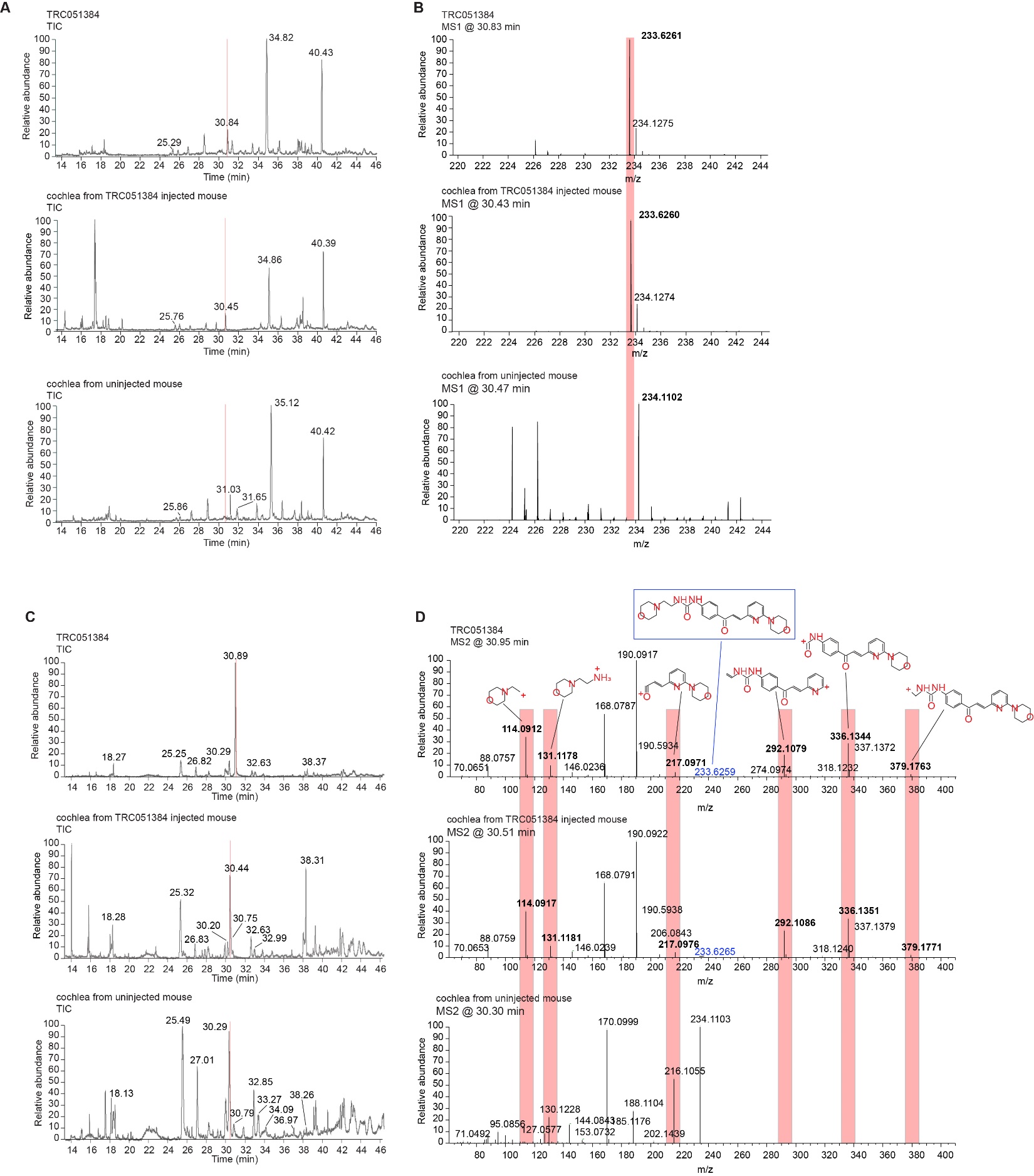

**Supplementary Figure 3. TRC051384 is readily detectable in cochlear extracts after intraperitoneal administration (60 mg/kg) by mass spectrometry (MS).**

**(A-B)** Panels show the MS total ion chromatogram (14–46 min) and the extracted MS1 spectra from the 220–244 *m/z* range at ~30 min (red highlight). The intact TRC051384 ion was consistently detected at ~30 minutes into a 60-minute LC–MS run in both the pure drug standard and cochlear extracts from treated mice. MS1 analysis further confirmed specificity, as the characteristic 233.62 *m/z* peak was absent in lysates from untreated controls but clearly present in both drug-alone and drug-treated samples.

**(C)** Targeted selected ion monitoring (tSIM) for the MS2 product ions of TRC051384 was again triggered at ~30 minutes into the LC–MS run, corresponding to the retention time of the precursor ion. This analysis provided additional confirmation of compound identity, as fragmentation yielded multiple diagnostic product ions characteristic of TRC051384, further validating its presence in the cochlear extracts.

**(D)** Product ions derived from TRC051384 were identified through fragment ion search (FISh) analysis in both the drug standard and cochlear extracts from treated mice. The FISh spectra revealed multiple diagnostic fragments that matched those expected from the ionization of TRC051384, thereby confirming its structural identity within the complex cochlear matrix (red highlights). The chemical structures and corresponding theoretical *m/z* values for each of the major product ions are listed alongside the observed spectra, providing additional validation of compound detection.

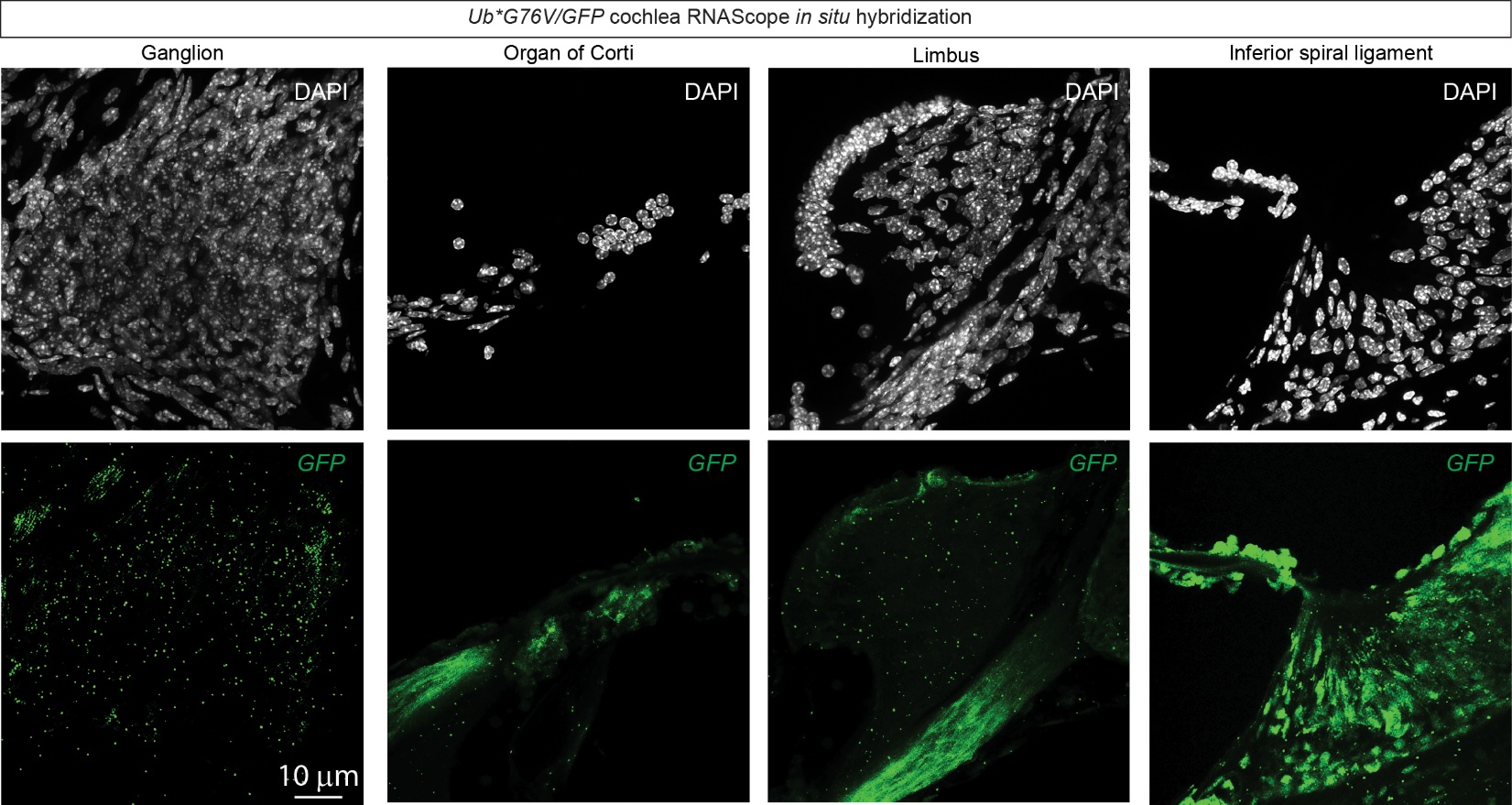

**Supplementary Figure 4. Widespread GFP mRNA expression in the cochlea of Tg(CAG-Ub*G76V/GFP)1Dant mice revealed by RNAscope *in situ* hybridization.**
Representative fluorescent RNAscope images of 12 µm mid-modiolar cochlear sections from P60 Tg(CAG-Ub*G76V/GFP)1Dant mice show robust GFP mRNA expression (green) across multiple cochlear regions, indicating broad transcriptional activity of the transgene.

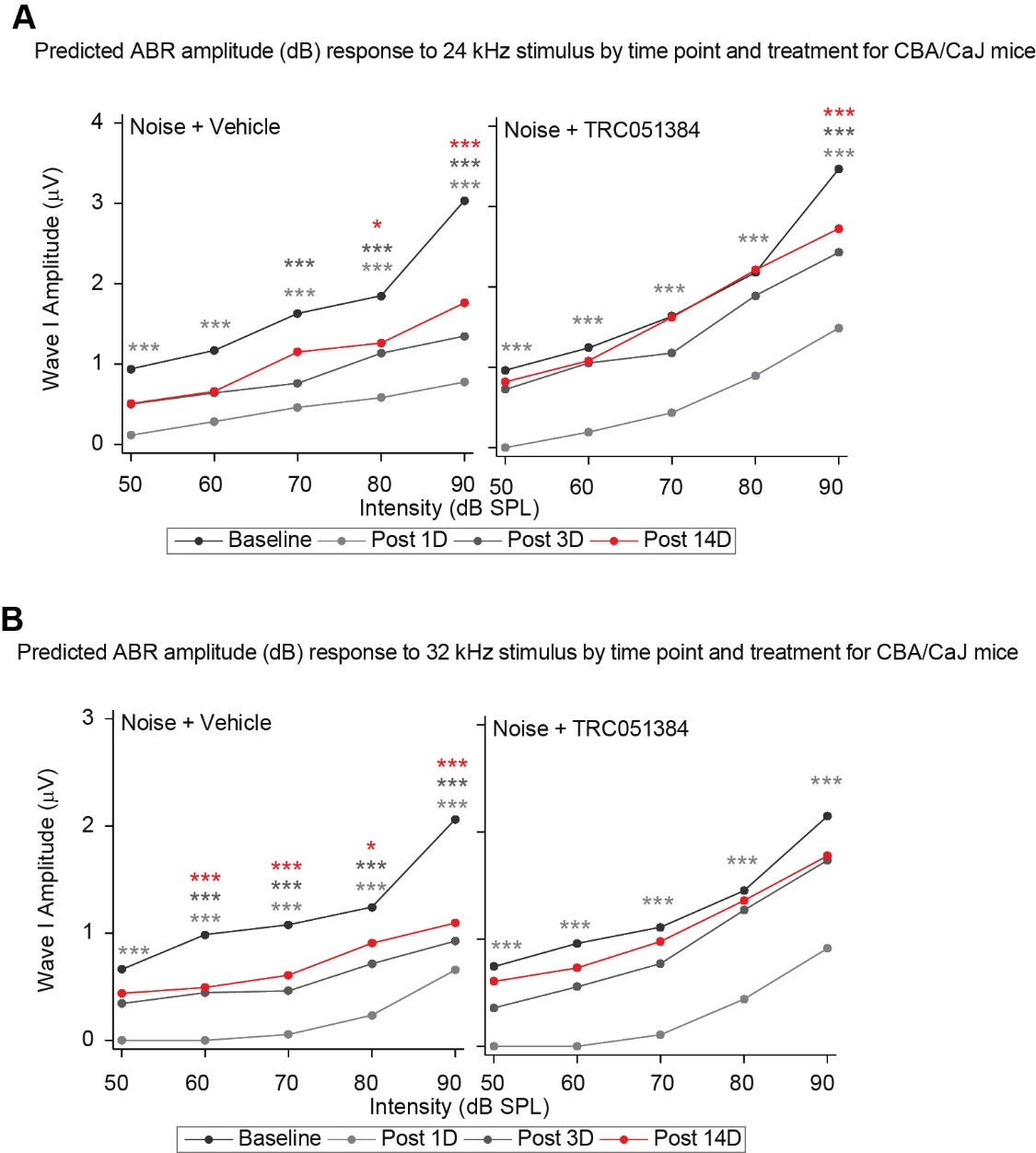

**Supplementary Figure 5. Independent confirmation that prophylactic TRC051384 treatment resulted in markedly greater Wave I amplitude recovery compared to vehicle controls.**

**(A)** Wave I amplitude failed to recover to BN levels by 14 DAN at 24 kHz in the vehicle treated group: 80 dB SPL = -0.59 µV adjusted 95% CI [-1.12, -0.052]), 90 dB SPL= -1.27 µV adjusted 95% CI [-1.81, -0.73]). TRC051384 treated mice failed to fully recover Wave I amplitudes to BN levels only at 90 dB SPL by 14 DAN ( -0.74 µV adjusted 95% CI [-1.24, -0.24]). A similar trend was observed at 3 DAN: the vehicle-treated group failed to recover at 70-, 80-, and 90-dB SPL, whereas the TRC051384-treated group exhibited partial recovery, failing only at 90 dB SPL.

**(B)** Wave I amplitude failed to recover to BN levels by 14 DAN at 32 kHz at intensities above 50 dB SPL in vehicle treated mice (60 dB SPL = -0.49 µV adjusted 95% CI [-0.83, -0.16]); 70 dB SPL= -0.47 µV adjusted 95% CI [-0.81, -0.13]); 80 dB SPL= -0.34 µV adjusted 95% CI [-0.67, -0.002]); 90 dB SPL= -0.96 µV adjusted 95% CI [-1.3, -0.63]). The TRC051384-treated group showed greater recovery. *** = p value < .001, ** < .01, * < .05 by one-way ANOVA determined significance with Tukey’s post hoc correction. (A-B) N = 6 male mice per group, (C) N = 8 male mice per group.

Supplemental Table Legends

**Table S1. Bulk RNA-seq analysis results.**
Differential gene expression analysis from bulk RNA sequencing corresponding to Figure 1H. The table includes normalized expression values for all genes passing the defined thresholds.

**Table S2. Functional enrichment (DAVID) analysis of Bulk RNA-seq results.**
Results of DAVID functional enrichment analysis performed on differentially expressed genes identified in Figure 1G. The table summarizes enriched Gene Ontology terms, KEGG pathways, enrichment scores, gene counts, and associated *p*-values or FDR and *p*-values.

**Table S3. GeoMx Digital Spatial Profiler (DSP) analysis.**
Summary of spatial transcriptomic profiling performed using the GeoMx Digital Spatial Profiler (DSP) for the regions shown in Figure 2H. The table includes region-of-interest (ROI) annotations, normalized expression values, and differential expression, and p values for transcripts measured across spatially resolved samples.

**Table S4. Functional enrichment (DAVID) analysis of GeoMx DSP data corresponding to Figure 2I.**
DAVID functional enrichment analysis of differentially expressed genes identified from the GeoMx DSP dataset presented in Figure 2I. The table summarizes enriched Gene Ontology terms, KEGG pathways, enrichment scores, gene counts, and associated *p*-values or FDR and *p*-values.
